# IL-12 restores the sequential cytotoxic capacities of anti-GD2 CAR-T and CAR-iNKT cells against glioblastoma

**DOI:** 10.64898/2026.08.23.746558

**Authors:** Tan-Dai Tran, Cindy Lamorlette, Loris Gerard, Jordan Brouard, Gianpietro Dotti, David Moulin, Loïc Reppel, Cécile Pochon, Marie-Thérèse Rubio

## Abstract

Glioblastoma (GBM) is a highly aggressive brain tumor characterized by rapid progression and a poor prognosis. CAR-based cellular therapies are promising approaches, and CAR-T cells targeting GD2 have demonstrated transient efficacy. Identifying how tumors evade these treatments is essential for advancing therapy development. In this study, we investigated the mechanisms through which GBM cells evade GD2.chimeric antigen receptor (CAR)-T and CAR-invariant natural killer T (iNKT) *in vitro* and explored ways to overcome tumor escape. GD2-targeted CAR-T and CAR-iNKT cells were tested in a stepwise in vitro model that repeatedly exposed them to GD2^+^ cell lines. While CAR effector cells effectively killed GD2^+^ GBM cells in short-term assays, their anti-tumor efficacy declined after repeated antigen exposures. Tumor escape mechanisms included reduced CAR expression, impaired proliferation, reduced production of cytokine, granzyme, and perforin, tumor downregulation of GD2, trogocytosis, and upregulation of the HLA-E/NKG2A inhibitory compared to MICA-B/NKG2D activation pathways on tumor and immune cells. Increasing effector cell numbers or adding IL-15 ± IL-7 partially improved CAR persistence but did not fully restored CAR effector functions. By contrast, IL-12 addition optimized tumor-killing capacity by increasing CAR effector cell proliferation, CAR surface expression, IFN-γ production and balancing HLA-E/NKG2A versus MICA-B/NKG2D pathways.

In conclusion, GD2.CAR-T and GD2.CAR-iNKT cells effectively target GBM but are susceptible to repeated antigen exposure, which IL-12 could counteract. These findings encourage further development of armored IL-12 CAR-T or CAR-iNKT cells and further investigation of the roles of HLA-E and MICA-B pathways in immunotherapy against GBM.

## Introduction

GBM accounts for nearly half of all malignant central nervous system (CNS) tumors, known for rapid progression and near-inevitable recurrence, due to its intrinsic biological aggressiveness and resistance[1, 2]. Despite aggressive multimodal treatment, median survival remains only 12-15 months, and five-year survival fall below 10%[1]. The current standard-of-care for GBM including maximal safe surgery followed by radiotherapy, temozolomide and chemotherapy modestly extend survival and fail to prevent disease recurrence[1–3]. Despite recent advances such as anti-angiogenic therapies[4], tumor-treating fields[5], and molecular targeted agents[6], durable tumor control remains unattainable.

Clinical trials with chimeric antigen receptor (CAR) T-cell therapy targeting GBM-associated antigens such as EGFRvIII, HER2, and IL-13R*a*2 have proven safe[7], but limited durable efficacy[8]. GD2, a disialoganglioside on glioma cells but limited in normal tissue, is an attractive cancer immunotherapy target[9]. Studies demonstrated that GD2.CAR-T cells mediated strong initial anti-tumor responses; however, sustained tumor control remains challenging[7, 10] due to CAR-T exhaustion[11–13] and the immunosuppressive GBM tumor microenvironment (TME)[14–16].

Recognizing conventional CAR-T therapy’s limitations, iNKT cells are emerging as a promising alternative immune effector. They are a unique T lymphocyte subset with an invariant T-cell receptor that recognizes glycolipid antigens via CD1d[17–19]. Bridging innate and adaptive immunity, iNKT cells respond rapidly by secreting proinflammatory cytokines such as IFN-γ, directly targeting tumor cells, and modulating immunosuppressive myeloid populations, particularly MDSCs and TAMs, within TME[18, 19], thereby reducing tumor immune evasion[20, 21]. iNKT cells display superior tumor-site trafficking, thereby having an advantage in TME[22]. CAR-iNKT cells combine tumor antigen specificity with innate immunomodulatory properties[23]. They show a lower risk of exhaustion and alloreactivity, allowing safer allogeneic use[18, 20]. In addition, neuroblastoma models show that GD2-targeted CAR-iNKT cells infiltrate tumors more efficiently than CAR-T cells[18, 20, 24, 25], but their effectiveness in GBM remains unexplored.

This study assessed the *in vitro* antitumor effects of anti-GD2 CAR-T and iNKT cells via serial re-exposure assays to understand functional changes in CAR effector cells after prolonged antigen exposure. The data described mechanisms associated with loss of CAR cell function and showed that IL-12 improves CAR-T and CAR-iNKT persistence and function across repeated GBM challenges.

## Materials and Methods

### Cell culture

Glioblastoma cell lines (LN229, T98G, U87) from CRAN laboratory (UMR7039, France) were cultured in RPMI-1640 GlutaMAX (Gibco) with 10% FBS (PAN Biotech) and 1% PS, incubated at 37°C with 5% CO₂. The immune cell medium consisted of RPMI 1640 Advanced (Gibco) with 8% FBS, 2 mM L-Glutamine (Sigma-Aldrich), and 1% PS.

### T and iNKT cell isolation and activation

Peripheral blood mononuclear cells (PBMCs) were obtained from Buffy coats (Etablissement Français du Sang Grand Est, France) under ethical guidelines and donor consent. PBMCs were isolated by density gradient centrifugation on lymphoseparation solution (Eurobio). T cells were purified by negative selection using the Pan T Cell Isolation Kit (Miltenyi Biotec). Invariant NKT cells were isolated from PBMCs by positive magnetic selection using anti-TCR Vα24-Jα18-PE staining (REA1054-6B11, Miltenyi Biotec), followed by anti-PE microbeads and MACS columns (Miltenyi Biotec). T and iNKT cells were activated for 3-5 days before transduction in completed RPMI-advanced with IL-15, IL-7 (10 ng/mL, PeroTech) and anti-CD3/CD-28 Dynabeads (1:1, Gibco) for T cells, and anti-CD3/anti-CD28 coated antibodies (MAB100/MAB342, 40 ng/mL, Bio-techne) and IL-15 (20 ng/mL) for iNKT cells.

### Vector construction

The GD2.CAR construct included a CD8α hinge/transmembrane domain, CD28-CD3ζ, and an IL-15 transgene, cloned into a γ-retroviral vector **(Fig.S1c-d).** This construct was kindly provided by Prof. Gianpietro Dotti (University of North Carolina).

### T and iNKT cell transduction

IL-15-secreting GD2-specific CAR-T cells were generated as previously described[26]. 4^th^ generation-retrovirus, T and iNKT cells (1x 10⁵ cells/well) were transduced with GD2.CAR construct on retronectin-coated plates (5 µg/cm²) preloaded with viral supernatants (spinoculation, 2000g, 90 min). These cells were then incubated in complete RPMI Advanced and expanded with cytokines; medium was renewed every 3-4 days until day 14, when both transduced and non-transduced cells were used for functional assays. The transduction rate was evaluated by flow cytometry on days 9-11 post-transduction.

3^rd^ generation-lentivirus, T cells (1x 10⁵ cells) were transduced with the GD2.CAR lentivirus containing two costimulatory domains (4.1.BB and CD28) with MOI 72 and protamine sulfate (100 *μg*/mL).

### Cytotoxicity assays

Short-term (16h) and long-term (72h) co-culture cytotoxicity assays evaluated the ability of CAR-T and CAR-iNKT cells to kill GD2^+^ GBM targets. GBM cell lines were seeded at 1×10⁵ cells per well in 24-well plates one day prior. The next day, effector cells at the indicated E:T ratios were added. For short-term assays with LN229, E:T ratios of 10:1, 5:1, 2:1, 1:1, 1:2, and 1:5 were tested. For long-term assays, immune and GBM cells were co-cultured at a 1:1 ratio. After incubation, cells were collected, stained with a viability dye, anti-CD3, and anti-B7H3 antibodies, and analyzed by flow cytometry to quantify residual tumor cells.

### Re-exposure challenge assays

Tumor antigen re-exposure assays evaluated effector cell persistence, cytotoxicity, and exhaustion, including non-transduced, CAR-T, and CAR-iNKT cells. LN229 and T98G were used for all challenges, while U87 and non-transduced immune cells were used only in the first cycle. GBM cells were seeded at 1×10^5^ cells per well (Corning) 7-24 hours before effector cell addition at a 1:1 (E:T) ratio in complete RPMI; some experiments used a 5:1 ratio with LN229. Co-cultures were maintained for 72 hours. After each cycle, non-adherent effector and adherent tumor cells were harvested for flow cytometry analysis. Some effector cells were re-seeded onto fresh GBM cells for the next challenge cycles. Additionally, some experiments included cytokines: IL-7 (10 ng/mL) + IL-15 (10 ng/mL) for CAR-T cells; IL-15 (20 ng/mL) for CAR-iNKT cells; and IL-12 (2 ng/mL) for both CAR-T and CAR-iNKT cells.

### Immunophenotyping

Flow cytometric analysis of surface and intracellular markers was performed using fluorochrome-conjugated antibodies (Sony, BD Biosciences, Cell Signaling Technology, BioLegend). Effector cells were stained for: CD3 (FITC, OKT3), CD4 (BV421, RPA-T4; PE/Cy7, SK3), CD8 (PerCP/Cy5.5, SK1), NKG2A (PE/Dazzle™ 594, S19004C), NKG2D (BV510, 1D11; PerCP 5.5, 1D11), GD2 (APC, 14G2a; BV421, 14.G2a), IFN-γ (BV421, 4S.B3), TNF-α (BV711, MAb11), granzyme B (PE/Dazzle™ 594, QA16A02), perforin (PE, dG9), LAG-3 (PE, 11C3C65), TIM-3 (PE/Dazzle™ 594, F38-2E2), PD-1 (BV510, EH12.2H7). Tumor cells were stained for GD2 (APC, 14G2a; BV421, 14.G2a), B7-H3 (BV421, 7-517), CD1d (PerCP/Cy5.5, 51.1), HLA-E (PE, 3D12), and MICA-B (APC, 6D4). CAR transduction efficiency and subset purity were confirmed using CD3 (FITC, OKT3), TCR Vα24 (PE, REA948-C15), and G4S–CAR linker (APC, E702V), with live/dead discrimination by APC-Cy7 Fixable Viability Stain 780 (BD). Data were acquired on BD Symphony and BD Celesta flow cytometers using FACSDiva and analyzed with FlowJo v10.

### Fluorescence microscopy

CAR effector and GBM cells were co-cultured for 4 hours in incubator. Coverslips, pre-coated with poly-L-lysine and incubated overnight at 4°C, were rinsed with PBS and air-dried before cell seeding.

Cells were harvested, washed twice with PBS, and stained for 20 minutes with anti-CD3 (FITC, OKT3, Sony) and anti-GD2 (BV421, 14.G2a, BD) antibodies. Stained cells were washed twice, fixed with 4% paraformaldehyde, and seeded onto poly-L-lysine–coated coverslips. Slides were stored in the dark at 4°C until imaging after adding Fluorescence mounting medium (Agilent, S302380-2). Fluorescence imaging was performed on a Zeiss AXIO fluorescence microscope using 40 and 63× oil-immersion objectives (Zeiss) under optimal imaging parameters. Images were processed and analyzed with ImageJ[27].

### Enzyme-linked immunosorbent assay (ELISA)

Culture media samples were used to quantify IFN-γ levels by ELISA using the DuoSet Human IFN-γ ELISA kit (DY285B, R&D Systems, USA). Briefly, 96-well plates were coated overnight at room temperature with the monoclonal capture antibody (2 µg/mL), then washed, blocked with reagent diluent (300 µL/well), and incubated with 1:100 diluted samples for 2 hours. Human IFN-γ detection antibody (200 ng/mL) was added for another 2 hours. Streptavidin–horseradish peroxidase conjugate and TMB substrate (100 µL/well) were added, the reaction was stopped with sulfuric acid (50 µL/well), and the optical density was measured at 450 nm. Samples were analyzed in duplicate by a microplate reader (Varioskan). IFN-γ concentrations were calculated from the curve and expressed as pg/mL.

### Statistical analysis

Data are presented as mean ± SEM (standard error of mean). Statistical analyses were conducted in GraphPad Prism (version 10, GraphPad Software, USA). Statistical result comparisons were performed between CAR-T and CAR-iNKT and among CAR-T and CAR-iNKT subgroups, including conditions with and without cytokine supplementation. Comparisons across or among different challenge cycles and with day 0 were performed using 2-way ANOVA followed by Turkey’s multiple comparisons test. Differences were considered statistically significant at *P*<0.05, *=p<0.05, **=*P*<0.01, ***=*P*<0.001, and ****=*P*<0.0001.

## Results

### Cytotoxicity of GD2.CAR-T and GD2.CAR-iNKT cells against GBM targets

Anti-GD2 CAR-T and CAR-iNKT cells were generated from highly purity sorted T and iNKT cells from healthy donor PBMCs (**Fig. 1a, Fig.S1a-b**). Using a 4^th^-generation anti-GD2 CAR (**Fig.S1c-d**), mean ± SEM CAR expression reached 54.23 ± 0.33% for CAR-T and 44.92 ± 3.28% for CAR-iNKT cells (n=24-27, p=0.036) (**Fig. 1a, Fig.S1e**). Starting from 10^5^ effector cells at day 0, CAR-T and CAR-iNKT expanded to 22.6 x 10^6^ ± 2.5 x 10^6^ cells (n=21) and 17.4 x 10^6^ ± 2.56 x 10^6^ cells (n=19), respectively, at day 14 (p=ns) (**Fig.1b**).

**Figure 1:**
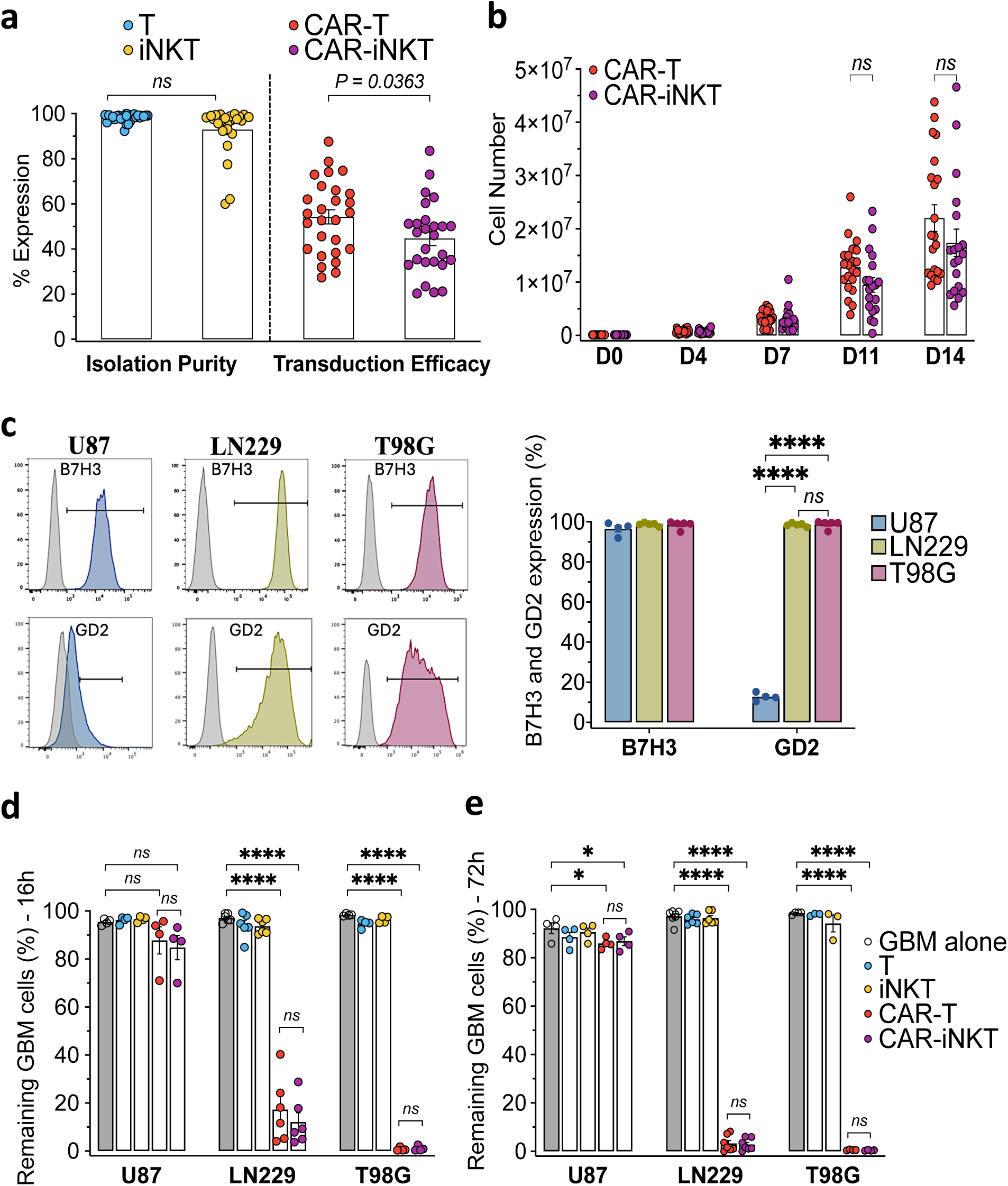
Generation of GD2.CAR-T and CAR-iNKT cells and assessment of their short-term cytotoxic activity against GBM cell lines. **a.** Percentage of T and iNKT cell purity after isolation and expansion (n=24-27) and proportions of GD2.CAR-expressing T and iNKT cells by day 11 after transduction. **b.** Numbers of GD2.CAR-T and CAR-iNKT cells before (D0) and up to Day 14 after transduction. **c.** Flow cytometry histograms showing surface expression of B7H3 and GD2 on 3 three glioblastoma cell lines: U87, LN229, and T98G. (n=4-5). **d-e.** Cytotoxic tests of T, iNKT, and GD2.CAR-T and CAR-iNKT cells against mentioned GBM cell lines performed in vitro at a ratio of 1:1. The percentage of remaining GBM cells (U87, LN229, and T98G) was measured by flow cytometry at 16 hours **(d)** and 72h **(e)** of co-culture. Comparisons performed with two-way ANOVA tests, ns = not significant, *=p<0.05, ****=p<0.0001.

We evaluated the direct cytotoxicity of transduced and untransduced T and iNKT cells against three glioblastoma cell lines: LN229 and T98G, which highly express GD2 (GD2^+^), and U87, expressing low GD2 levels (GD2^low^) (**Fig.1c**). All three tumor cell lines showed high B7H3 expression, enabling the identification of residual tumor cells in cytotoxicity assays (**Fig.1c, Fig.S2a**). We first assessed the cytotoxic activity of GD2.CAR-T and GD2.CAR-iNKT cells against LN229 cells after 16-hour co-culture at increasing E:T ratios (**Fig.S2b**). Both CAR effector populations displayed comparable dose-dependent killing, with complete LN229 elimination at 10:1 and 5:1 ratios and approximately 80% tumor-cell elimination at a 1:1 ratio (**Fig.S2c**). This effect was GD2-dependent, as untransduced T and iNKT cells showed no cytotoxic activity (**Fig.S2c, Fig.1d-e**). Additional 1:1 cytotoxicity assays across the three GBM cell lines showed that both GD2.CAR-T and GD2.CAR-iNKT cells efficiently and similarly killed GD2^+^ targets, but not GD2^low^ U87 cells, in both short-term (16h) and long-term (72h) assays (**Fig.1d-e**).

### Progressive loss of cytotoxicity during GBM re-exposure assays due to GD2.CAR effector dysfunctions

We next performed serial re-exposure of GD2.CAR effectors to GD2^+^ GBM cell lines to assess their functional capacity to respond to sequential tumor stimulations. In these assays, LN229 and T98G were seeded before adding CAR effectors at 1:1 E:T ratio (**Fig.S3a**). At the end of each 3-day cycle, whole cells were collected for phenotype characterization and functional assays.

We observed that both GD2.CAR-T and GD2.CAR-iNKT cells progressively lost their ability to control GBM cell lines *in vitro* after repeated antigen stimulation (**Fig.2a**). CAR-T cells maintained strong cytotoxicity against LN229 and T98G during the first two cycles. CAR-iNKT cells showed similar activity against T98G, whereas their killing of LN229 began to decline from cycle 2. By cycle 3, both GD2.CAR-T and GD2.CAR-iNKT cells displayed markedly impaired control of both GBM cell lines (**Fig.2a**).

**Figure 2:**
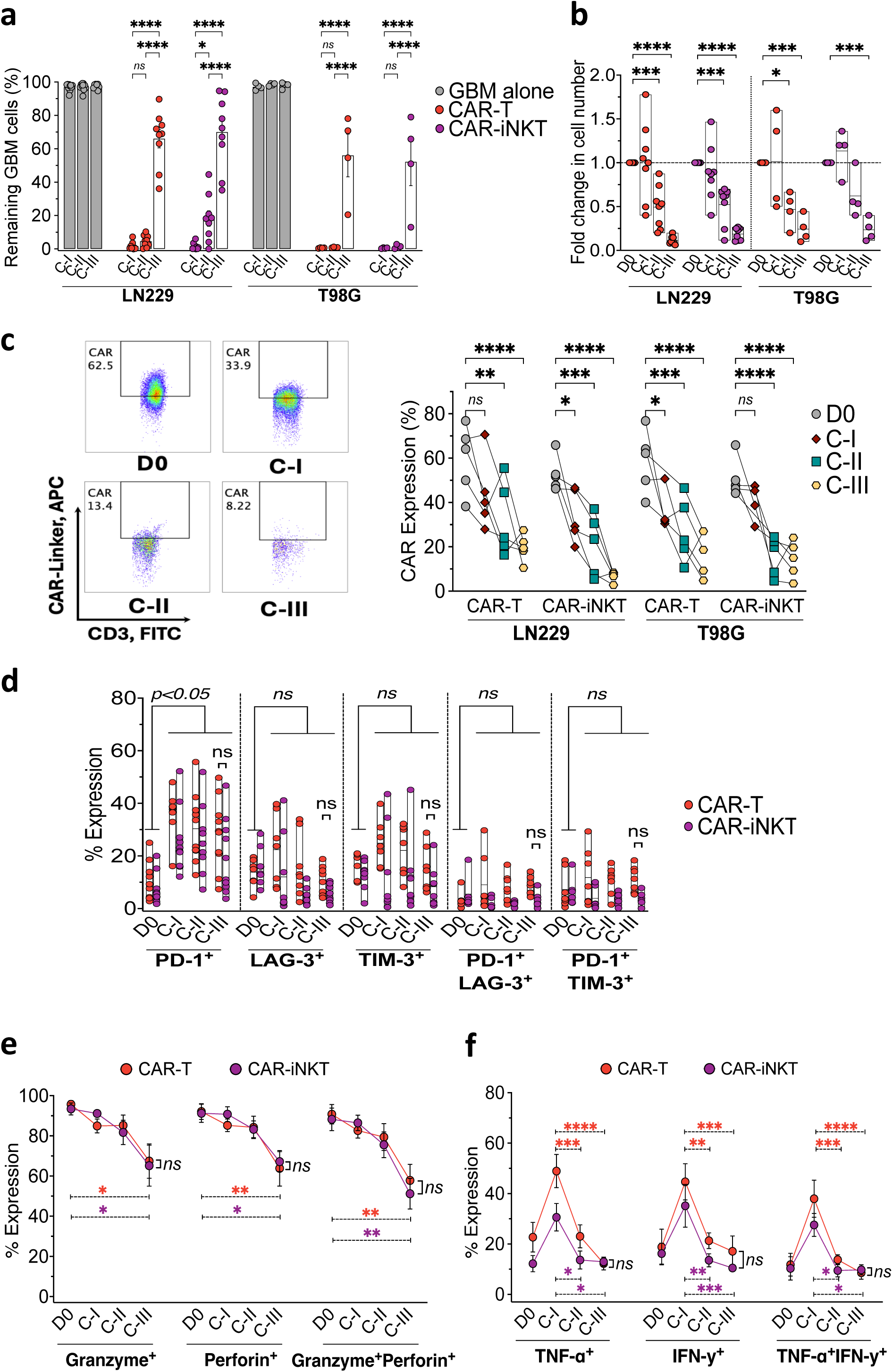
GD2.CAR-T and CAR-iNKT cells progressively lose cytotoxic activity against GBM cells during sequential tumor challenge assays. **a.** Percentage of surviving LN229 (n=9) and T98G (n=4) cells after co-culture with GD2.CAR-T (red) or CAR-iNKT (purple) cells across 3 tumor challenge cycles (C-I, C-II, and C-III). While both CAR-T and CAR-iNKT cells showed strong cytotoxicity at cycle 1, a reduction in cytotoxicity in later cycles was observed with significantly increased GBM remaining cells compared to cycle 1 (p<0.0001). **b.** Fold change of total effector cell numbers relative to day 0 (D0) across tumor challenge cycles with LN229 (n=9) and T98G (n=4). **c. Left panel:** flow cytometry dot blot illustrating GD2.CAR surface expression (anti-G4S linker monoclonal antibody) detection on CD3^+^ GD2^-^ viable effector cells on D0 and after each LN229 challenge cycle. **Right panel:** percentages of CD3^+^ effector cells expressing GD2.CAR from D0 to cycle 3 against LN229 (n=5). **d.** PD-1, LAG-3, TIM-3 expression alone or in combination on GD2.CAR-T (n=5, red) and CAR-iNKT (n=5, purple) cells on D0 and after LN229 challenge cycles. **e-f.** Intracellular granzyme ± perforin (**e**) or TNF-α ± IFN-γ (**f**) expression on GD2.CAR-T (n=6, red) and CAR-iNKT (n=6, purple) cells on D0 and after LN229 challenge cycles. Comparisons performed using two-way ANOVA, Tukey’s multiple comparisons tests., ns = not significant, *=p<0.05, **=p<0.01, ***=p<0.001, ****=p<0.0001.

The loss of cytotoxic capacity against both GD2^+^GBM cell lines after cycle 3 was associated with significant drop of effector cell numbers throughout the cycles (**Fig.2b**), as well as of loss of CAR molecule expression on CAR effectors’ surface notably after cycles 2 and 3 (**Fig.2c**) leading to marked declined of absolute numbers of GD2.CAR-T and CAR-iNKT cells against LN229 and T98G (**Fig.S3b**).

Analysis of T-cell activation and exhaustion markers showed that both GD2.CAR effector populations progressively upregulated PD-1 from cycle 1 to cycle 3 compared with D0 (**Fig.2d**). In contrast, LAG-3 and TIM-3 did not increase over the cycles relative to D0. Co-expression of PD-1⁺LAG-3⁺ and PD-1⁺TIM-3⁺ remained low and stable, with no significant variation across cycles (**Fig.2d**). Despite displaying an activation rather than an exhaustion phenotype, GD2.CAR effectors showed reduced granzyme and/or perforin production by cycle 3 (**Fig.2e, Fig.S3c**) and progressive decline of TNF-α and/or IFN-γ production from cycle 1 to cycle 3 (**Fig.2f, Fig.S3d**).

### Tumor cell alteration during immune pressure by GD2.CAR effectors

We then investigated whether tumor escape during repeated challenges could be driven by GBM adaptation to CAR-effector pressure. We first assessed HLA-E and MICA-B, the respective ligands of the inhibitory NKG2A and activating NKG2D receptors[28, 29], on residual GD2^+^ GBM cells after the third challenge. Although LN229 and T98G cells did not express these ligands before immune exposure, both cell lines showed marked upregulation of the two ligands after GD2.CAR-T and CAR-iNKT contact (**Fig.3a-c**). By cycle 3, HLA-E upregulation exceeded that of MICA-B, increasing the HLA-E/MICA-B ratio during CAR-effector exposure. This increase was particularly significant in both GBM cell lines exposed to CAR-iNKT cells (**Fig.3d**). In parallel, we monitored NKG2A and NKG2D expression on GD2.CAR effectors across tumor challenge cycles (**Fig.3e, Fig.S3e**) and observed a significant increase of the NKG2A/NKG2D ratio on CAR-iNKT cells exposed to LN229 (**Fig.3f**). No significant change was observed after exposure to T98G (**Fig.S3f**). These results suggest that the HLA-E/NKG2A inhibitory axis may contribute to tumor escape, particularly in GD2.CAR-iNKT cells challenged with LN229.

**Figure 3:**
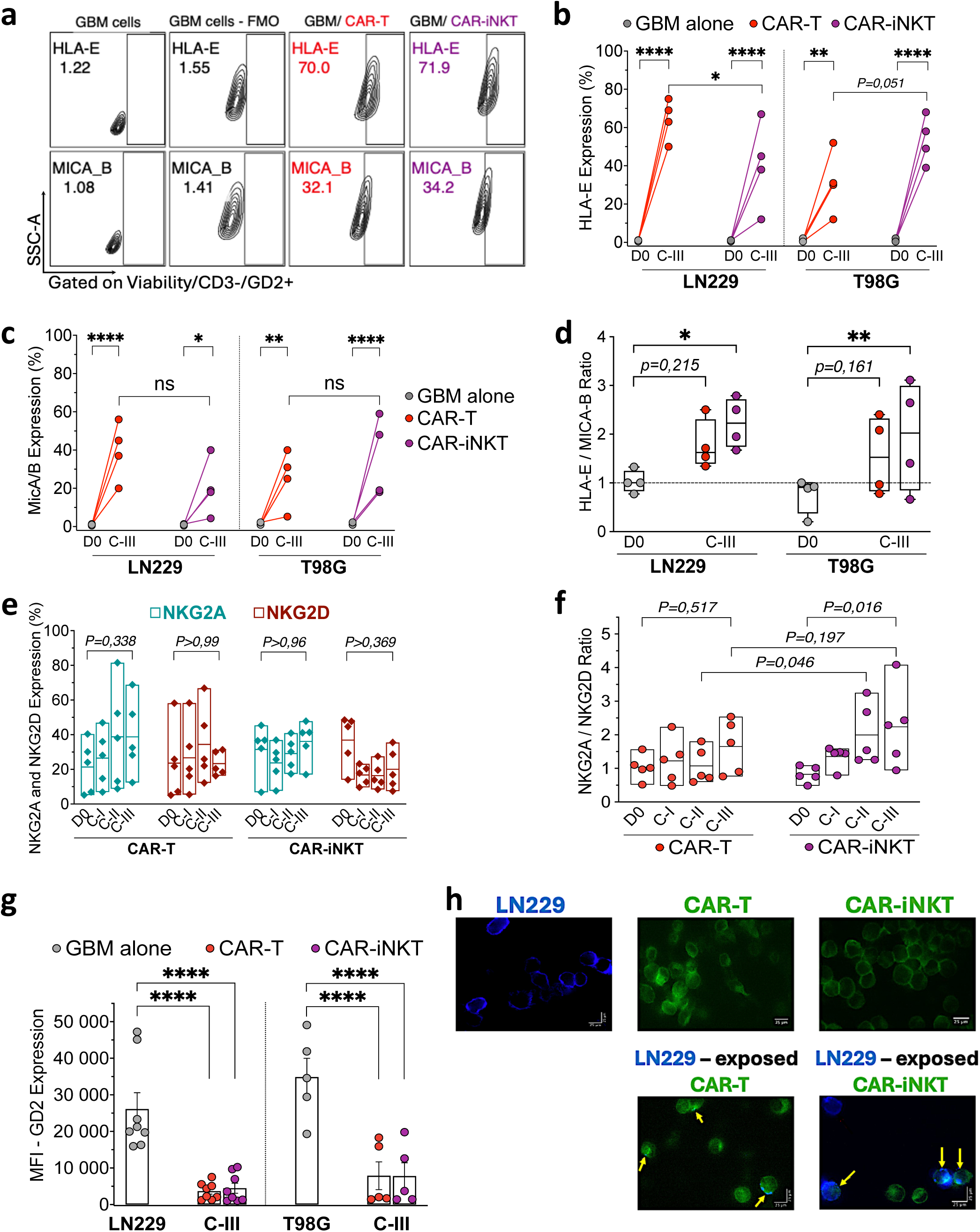
Exploration of NK inhibitory/activation pathways, antigen loss and trogocytosis during sequential tumor challenges. **a.** Flow cytometry plot showing the expression of HLA-E and MICA-B on remaining GBM viable cells after cycle 3 of co-culture with GD2.CAR-T (red) and CAR-iNKT (purple) cells. **b-c.** Proportions of HLA-E **(b)** and MICA-B **(c)** expressing LN229 (n=4) and T98G (n=4) cells after cycle 3 of co-culture with GD2.CAR-T (red) and CAR-iNKT (purple) cells. **d.** HLA-E/MICA-B ratio measured on GD2^+^ cell lines GBM cells on day 0 (D0) and after contact with GD2.CAR-T (red) and CAR-iNKT (purple) cells at cycle 3. **e.** Proportions of GD2-CAR-T (n=4) and CAR-iNKT (n=4) expressing the NK inhibitory receptor NKG2A (turquoise) and the activating receptor NKG2D (brown) at D0 and after each tumor challenge cycle. **f.** NKG2A/NKG2D ratio on GD2.CAR-T (n=4, red) and CAR-iNKT (n=4, purple) cells at D0 and after each tumor challenge cycle. **g.** Median florescence intensity (MFI) of GD2 expression on GD2^+^ GBM cells before (n=5-8, grey) and after cycle 3 of co-culture with GD2.CAR-T (n=5-8, red) and CAR-iNKT (n=5-8, purple). **h. Upper panels:** Fluorescence microscopy images (40×objective) showing GD2 antigen staining on LN229 cells (anti-GD2-BV421, blue) and CD3 staining on CAR-T and CAR-iNKT cells (anti-CD3-FITC, green). **Lower panels:** the same staining performed on GD2.CAR-T and CAR-iNKT cells collected after 4 hours of contact with LN229 cells. Yellow arrows indicate GD2 signal detection (blue spots) on the surface of GD2.CAR effector cells (green cells). Comparisons performed with Two-way ANOVA, Tukey’s multiple comparisons test. ns = not significant, *=p<0.05, **=p<0.01, ****=p<0.0001.

Antigen loss is a common tumor escape mechanism to CAR-T cells[30, 31]. We therefore analyzed the intensity of GD2 expression on GBM cells after 72h of exposure to CAR effector cells at the end of cycle 3. In comparison to non-exposed LN229 and T98G cells to GD2.CAR effectors, GBM cells significantly downregulated GD2 expression intensity under GD2.CAR effector pressure (**Fig.3g**). Furthermore, in parallel with GD2 down expression in GBM cell lines, we observed that GD2 antigen was detectable by fluorescence microscopy on both GD2.CAR-T and CAR-iNKT cell surfaces after 4h co-culture with GD2^+^GBM cells suggesting trogocytosis (**Fig.3h**), a phenomenon involving the transfer of membrane fragments between cells identified as a mechanism by which CAR-T cells uptake tumor antigens[30, 31]. We confirmed the expression of GD2 on GD2.CAR effectors by flow cytometry with highest levels of expression at cycle 1 (**Fig.S3g**).

Together, these findings show that repeated exposure to tumor antigen drives CAR-effector dysfunction, characterized by reduced proliferation, impaired cytokine and cytotoxic granule production, and trogocytosis-associated tumor escape through antigen loss. For LN229 cells, this escape was also linked to a shift in the balance between NK activating and inhibitory pathways.

### Higher CAR cell input or cytokine addition with IL-15 ± IL-7 improved tumor control without reversing CAR dysfunction

We next tested whether increasing the number of GD2.CAR effector cells could improve their efficacy across re-exposure cycles. We first observed that increasing the E:T to 5:1 could partially improve tumor control after cycle 3 thanks to increase numbers of effector cells by cycle 3 but GD2.CAR effector functions still declined over the cycles (**Fig.S4**).

We next examined whether cytokine supplementation could support CAR-effector proliferation and functionality during repeated tumor challenges. Adding IL-7/IL-15 to GD2.CAR-T cells or IL-15 to CAR-iNKT cells (**Fig.S5a**) significantly improved control of both GD2^+^ GBM cell lines by cycle 3 compared with cytokine-free conditions (**Fig.4a-b**), whereas α-GalCer alone did not improve CAR-iNKT activity (data not shown). Cytokine supplementation preserved total effector-cell and GD2.CAR-effector viability across the three tumor challenge cycles (**Fig.4c-d, Fig.S5b**). However, it did not prevent GD2.CAR loss from the effector-cell surface (**Fig.4e-f**) or the progressive decline in IFN-γ, TNF-α, granzyme and perforin production against LN229 or T98G cells (**Fig.4g-h, Fig.S5c-f**). Activation and exhaustion marker expression on GD2.CAR effectors was comparable across cycles with or without cytokines, except for higher TIM-3 expression in CAR-T cells and higher PD-1 expression in CAR-iNKT cells, while PD-1^+^TIM-3^+^ and PD-1^+^LAG-3^+^ double-positive populations remained low (**Fig.S6a-b**).

**Figure 4:**
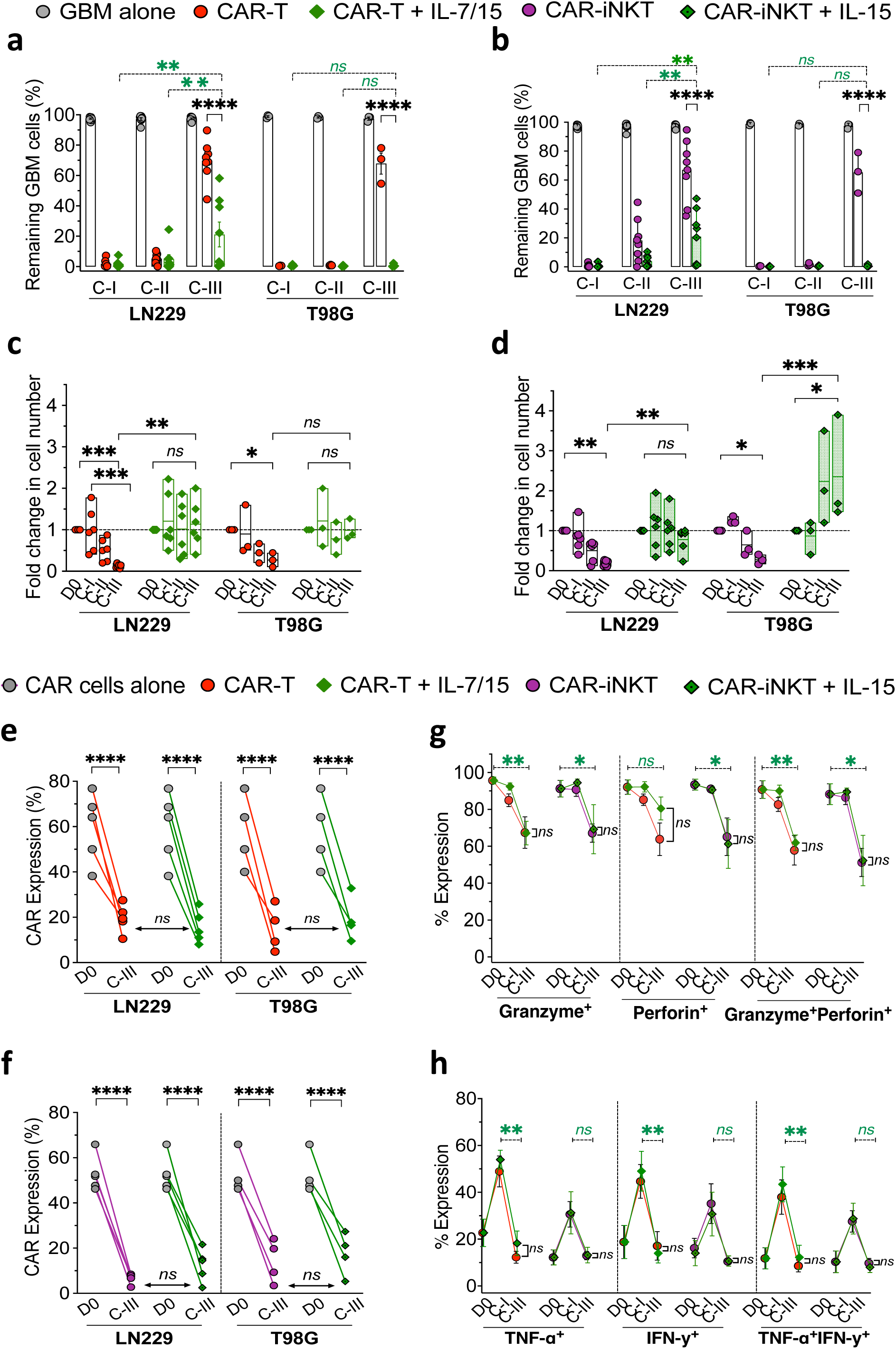
Impact of IL-15 ± IL-7 supplementation on cytotoxicity, CAR-expression, and cytokine and cytolytic molecule expression in CAR effectors. a-b. Percentage of residual LN229 (n=5) and T98G (n=3) cells after co-culture with GD2.CAR-T alone (red) or with IL7/IL15 supplementation (light green) (**a**) and with GD2.CAR-iNKT alone (purple) or with IL-15 supplementation (dark green) (**b**) after each tumor challenge cycles. **c-d.** Fold change of total effector cell numbers relative to day 0 (D0) across tumor challenge cycles with LN229 (n=6) and T98G (n=3) for GD2.CAR-T alone (red) or with IL7/IL15 supplementation (light green) (**c**), and for GD2.CAR-iNKT alone (purple) or with IL- 15 supplementation (dark green) (**d**). **e-f.** Percentages of CD3^+^ effector cells expressing GD2.CAR at D0 and after cycle 3 against LN229 (n=5) and T98G (n=3) for GD2.CAR-T alone (red) or with IL7/IL15 supplementation (light green) (**e**), and for GD2.CAR-iNKT alone (purple) or with IL-15 supplementation (dark green) (**f**). **g-h.** Intracellular granzyme ± perforin (**g**) and TNF-α ± IFN-γ (**h**) expression on GD2.CAR-T alone (red) or with IL7/IL15 supplementation (light green), and on GD2.CAR-iNKT alone (purple) or with IL-15 supplementation (dark green) on D0 and after cycle 1 and 3 of co-culture with LN229 cells (n=5). Comparisons performed with Two-way ANOVA, Tukey’s multiple comparisons test. Ns = not significant, *=p<0.05, **=p<0.01, ***=p<0.001, ****=p<0.0001.

Cytokine supplementation appeared to increase both MICA-B and HLA-E expression on LN229 cells in similar proportions, thereby maintaining a balanced HLA-E/MICA-B ratio (**Fig.5a-c**). Similarly, it enhanced the expression of both inhibitory and activating NK receptors, resulting in a balanced NKG2A/NKG2D ratio on GD2.CAR-T and GD2.CAR-iNKT effectors after repeated LN229 exposure (**Fig.5d-e, Fig.S6c-e**). Finally, cytokine supplementation did not avoid trogocytosis (**Fig.5f-g**) and GD2 antigen loss (**Fig.S6f**).

**Figure 5:**
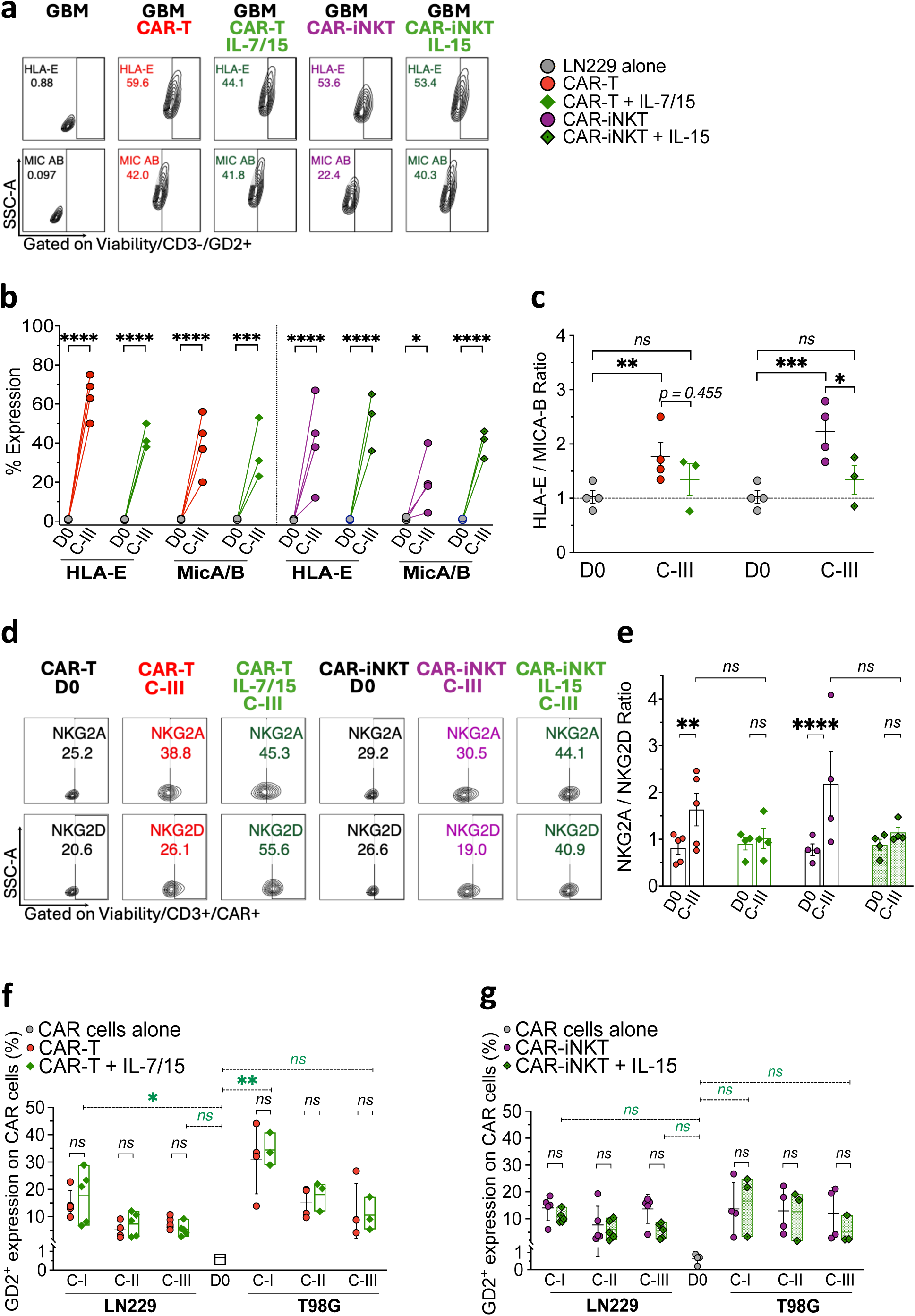
Impact of IL15 ± IL7 supplementation on NK inhibitory/activation pathways and trogocytosis during sequential tumor challenges a. Flow cytometry plot showing the expression of HLA-E and MICA-B on remaining viable LN229 cells after cycle 3 of co-culture with GD2.CAR-T alone (red) or with IL7/IL15 supplementation (light green), and with GD2.CAR-iNKT alone (purple) or with IL-15 supplementation (dark green). **b.** Proportions of HLA-E- and MICA-B-expressing LN229 cells (n=4) at day 0 (D0) after cycle 3 of co-culture with GD2.CAR-T alone (red) or with IL7/IL15 supplementation (light green), and with GD2.CAR-iNKT alone (purple) or with IL-15 supplementation (dark green). **c.** HLA-E/MICA-B ratio measured on LN229 cells on D0 and after contact with GD2.CAR-T alone (red) or with IL7/IL15 supplementation (light green), and with GD2.CAR-iNKT alone (purple) or with IL-15 supplementation (dark green) at cycle 3. **d.** Flow cytometry plot showing the expression NKG2A and NKG2D at D0 and after the third LN229 challenge cycle on GD2.CAR-T alone (red) or with IL7/IL15 supplementation (light green), and on GD2.CAR-iNKT alone (purple) or with IL-15 supplementation (dark green). **e.** NKG2A/NKG2D ratio on GD2.CAR-T alone (red) or with IL7/IL15 supplementation (light green), and on GD2.CAR-iNKT (purple) or with IL-15 supplementation (dark green) cells at D0 and after the third LN229 challenge (n=4 for each). **f-g.** GD2 antigen expression on GD2.CAR-T alone (red) or with IL7/IL15 supplementation (light green) (**f**), and on GD2.CAR-iNKT (purple) or with IL-15 supplementation (dark green) cells (**g**) after each tumor challenge with GBM cell lines (n=4 for each). Comparisons performed with Two-way ANOVA, Tukey’s multiple comparisons test. ns = not significant, *=p<0.05, **=p<0.01, ***=p<0.001, ****=p<0.0001.

Overall, IL-7/IL-15 supplementation during tumor challenges improved GBM control mainly by increasing GD2.CAR-T and CAR-iNKT cell numbers and by limiting the NKG2A inhibitory pathway against LN229 cells. Besides, CAR-effectors still become dysfunctional after cycle 3.

### IL-12 supplementation fully rescued GD2.CAR-T and CAR-iNKT function during the GBM challenge

IL-12 has been reported to enhance the persistence of CAR-T and CAR-iNKT cells, especially anti-GD2 CAR-iNKT cells in neuroblastoma models [32, 33]. We therefore explored whether IL-12 addition could sustain the cytotoxic function of CAR effectors in sequential exposure to GD2^+^GBM cells (**Fig.S7a**). We observed that IL-12 fully restored the cytotoxicity of both GD2.CAR-T and CAR-iNKT cells against LN229 and T98G up to 3 tumor challenges **(Fig.6a-b**). Addition of IL-12 helped to maintain the number of effector cells (**Fig.6c-d**), avoid the loss of the CAR molecules on GD2.CAR-T and CAR-iNKT cells (**Fig.6e-f, Fig.S7b-d**) and was associated with a significant expansion of both CAR effectors during the 3 tumor challenge cycles (**Fig.S7e**). Activation and exhaustion marker expression were significantly upregulated on GD2. CAR-T and CAR-iNKT cells in the presence of IL-12 compared to CAR effectors without IL-12 support, including increased proportions of PD-1⁺LAG-3⁺ and PD-1⁺TIM-3⁺, more notably on CAR-iNKT cells, suggesting higher rate of exhaustion, despite improved cytotoxic capacities by cycle 3 (**Fig.6g-h**). Additionally, IL-12 allows the maintenance of high and stable levels of IFN-γ, TNF-α, granzyme and perforin production by both GD2.CAR effectors in response to LN229 (**Fig.7a-b)** and T98G (**Fig.S8a-b)** GBM cell lines. As a result, culture media IFN-γ levels were significantly increased by the addition of IL-12 at each challenge cycles with both GD2.CAR effectors (**Fig.7c**). NKG2D was increased on both CAR-T and CAR-iNKT cells after contact with LN229 cells and NKG2A/NKG2D ration remained balanced across the cycles, in contrast with an increase of the inhibitory pathway in the absence of IL-12 (**Fig.7d-e, Fig.S8c-e**). Intriguingly, the addition of IL-12 was associated with significant GD2 expression on both GD2.CAR-T and CAR-iNKT effector cells after co-culture with LN229 and T98G cells (**Fig.7f**), together with persistent trogocytosis in the presence of IL-12 (**Fig.S9**). These findings suggest that trogocytosis may serve as a surrogate marker of CAR-effector cytotoxic activity. The beneficial effect of IL-12 was further confirmed in CAR-T cells expressing a third-generation anti-GD2 CAR delivered by a lentiviral vector, where IL-12 prolonged LN229 tumor control for up to seven challenge cycles (**Fig.S10**).

**Figure 6:**
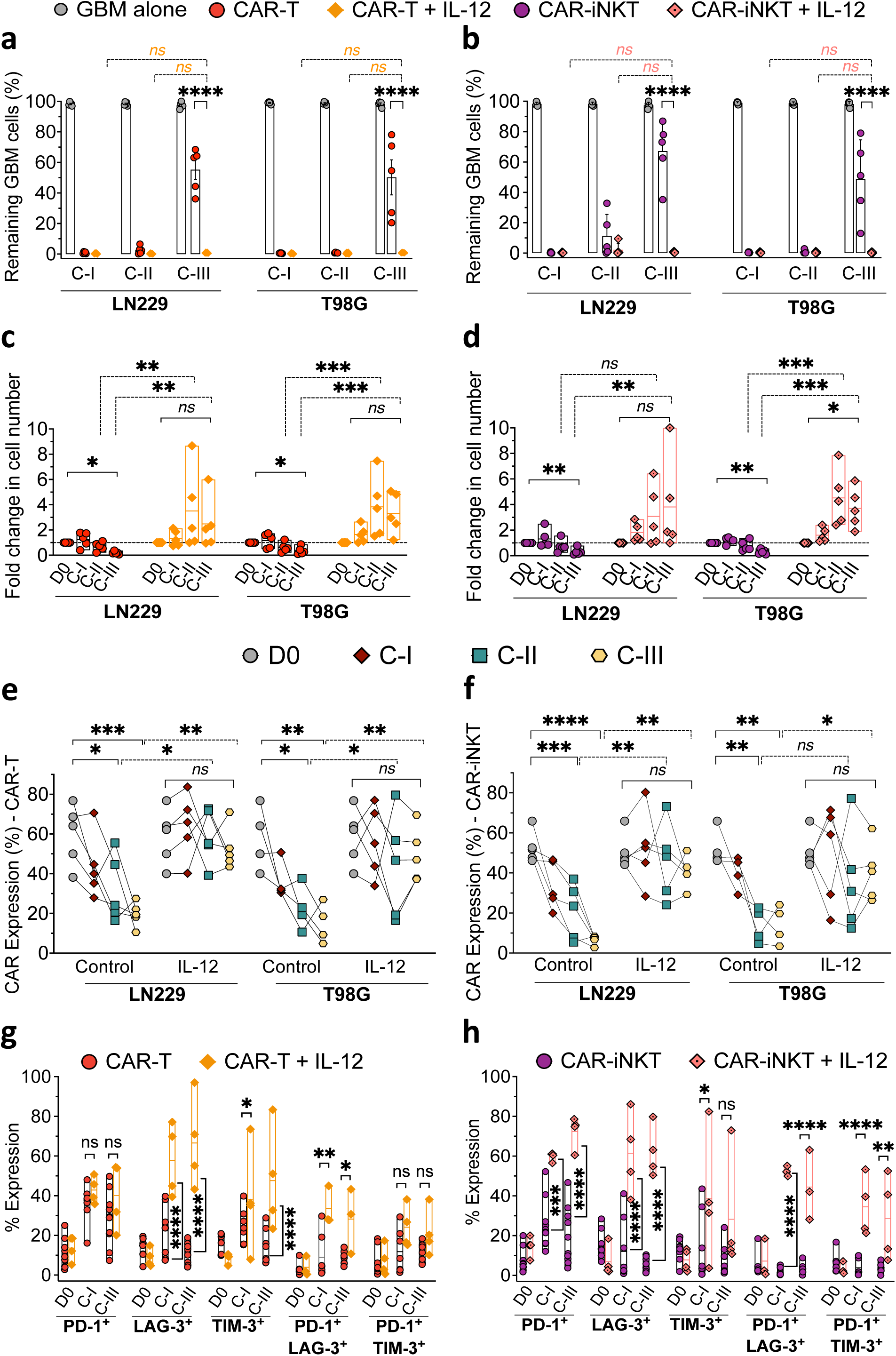
IL-12 restores GD2.CAR-T and GD2.CAR-iNKT sequential cytotoxicity by maintaining GD2.CAR-expression and enhancing effector expansion capacities and activation. a-b. Percentage of residual LN229 (n=5) and T98G (n=5) cells after co-culture with GD2.CAR-T alone (red) or with IL12 (orange) (**a**) and with GD2.CAR-iNKT alone (purple) or with IL-12 (pink) (**b**) after each tumor challenge cycles. **c-d.** Fold change of total effector cell numbers relative to day 0 (D0) across tumor challenge cycles with LN229 (n=5) and T98G (n=5) for GD2.CAR-T alone (red) or with IL12 (orange) (**c**) and for GD2.CAR-iNKT alone (purple) or with IL-12 (pink) (**d**). **e-f.** Percentages of CD3^+^ effector cells expressing GD2.CAR at D0 (grey) and after each tumor challenge cycle LN229 (n=5) and T98G (n=5) for GD2.CAR-T alone (red) or with IL12 (orange) (**e**) and for GD2.CAR-iNKT alone (purple) or with IL-12 (pink) (**f**). **g-h.** Proportions of PD-1, LAG-3, TIM-3 expression alone or in combination on GD2.CAR-T alone (red) or with IL12 (orange) (**g**), and on GD2.CAR-iNKT alone (purple) or with IL-12 (pink) (**h**) at D0 and after cycle 1 and 3 of co-culture with LN229 cells (n=5). Comparisons performed with Two-way ANOVA, Tukey’s multiple comparisons test. ns = not significant, *=p<0.05, **=p<0.01, ***=p<0.001, ****=p<0.0001.

**Figure 7:**
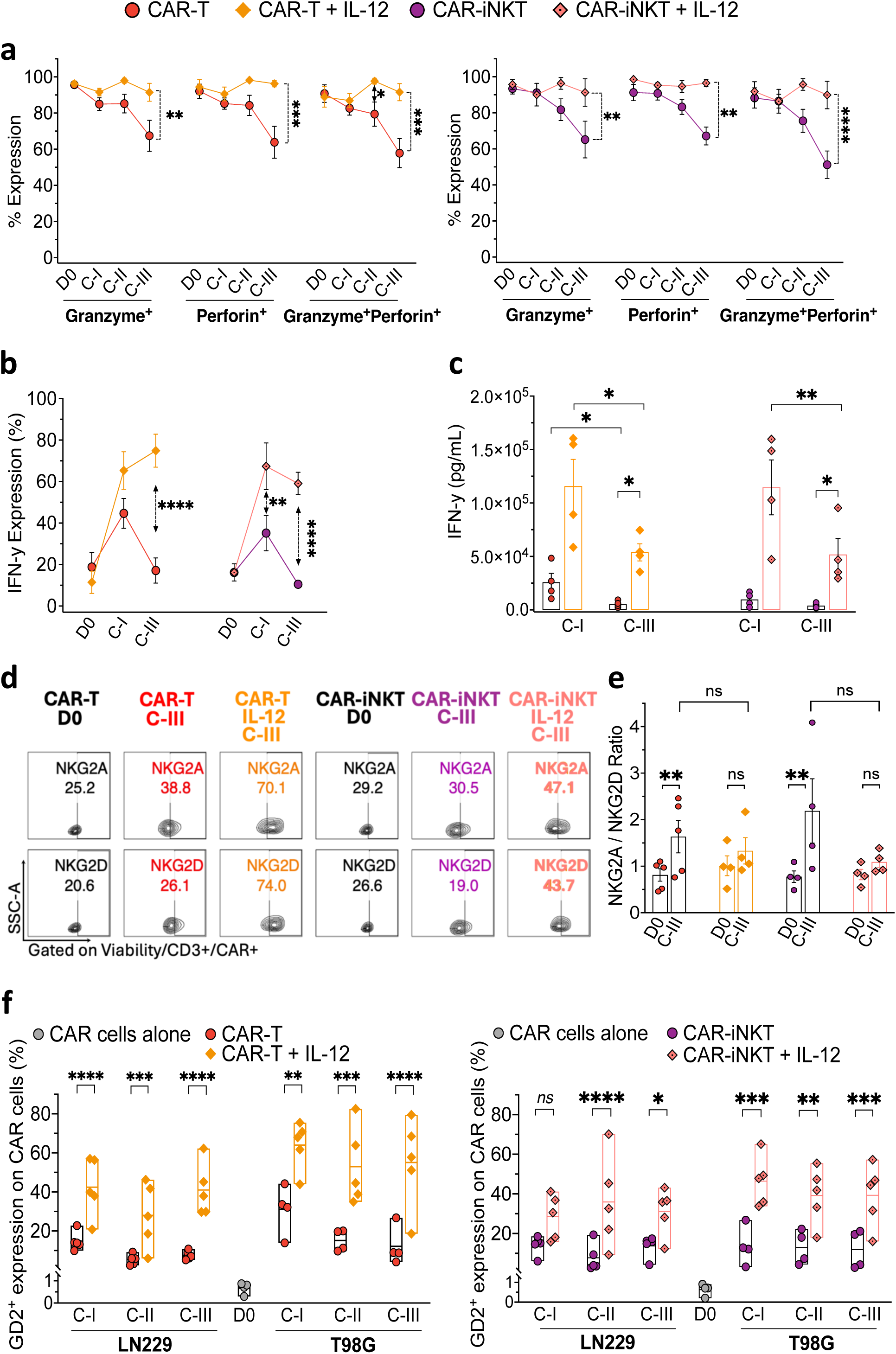
IL-12 maintains IFN-γ producing highly cytotoxic GD2.CAR-T and GD2.CAR-iNKT cells. **a.** Intracellular granzyme ± perforin expression on GD2.CAR-T alone (red) or with IL12 (orange) (**left panel**), and on GD2.CAR-iNKT alone (purple) or with IL-12 (pink) (**right panel**) on day 0 (D0) and after each LN229 challenge cycle (n=5). **b.** Intracellular IFN-γ expression on GD2.CAR-T alone (red) or with IL12 (orange), and on GD2.CAR-iNKT alone (purple) or with IL-12 (pink) on D0 and after cycle 1 and 3 of LN229 challenge (n=5). **c.** IFN-γ concentrations in culture media at the end of cycle 1 and 3 of LN229 challenge in the presence of GD2.CAR-T alone (red) or with IL12 (orange), and of GD2.CAR-iNKT alone (purple) or with IL-12 (pink) (n=5). **d-e.** Flow cytometry plot showing the expression the NK inhibitory receptor NKG2A and the activating receptor NKG2D (**d**) and NKG2A/NKG2D ratio (**e**) on GD2.CAR-T alone (red) or with IL12 (orange), and on GD2.CAR-iNKT alone (purple) or with IL-12 (pink) at D0 and after the third LN229 challenge cycle (n=5). **f**. GD2 antigen expression on GD2.CAR-T alone (red) or with IL12 (orange) (**left panel**), and on GD2.CAR-iNKT alone (purple) or with IL-12 (pink) (**right panel**) after each tumor challenge with GBM cell lines (n=5 for each condition). Comparisons performed using two-way ANOVA, Tukey’s multiple comparisons tests., ns = not significant, *=p<0.05, **=p<0.01, ***=p<0.001, ****=p<0.0001.

Overall, these findings indicate that IL-12 preserves CAR-T and CAR-iNKT functional persistence across repeated tumor challenges by maintaining CAR expression, sustaining a highly cytotoxic capacities, and promoting CAR-effector proliferation.

## Discussion

Despite encouraging response rates reported with CAR-T cells in early-phase clinical studies for GBM, most patients relapse[34]. We used *in vitro* assays to investigate how GD2-expressing GBM cells escape from GD2.CAR-T and CAR-iNKT cells. Our results show that both CAR-T and CAR-iNKT cells initially control GBM cells but progressively lose their antitumor activity after repeated tumor stimulation. This decline is linked to CAR-effector dysfunction and tumor antigen downregulation, consistent with recent clinical observations that EGFR-IL13Rα2-CAR-T cells expand in GBM patients with an activated phenotype during the first week but become exhausted by day 21, with reduced proliferative capacity, while residual tumor cells downregulate EGFR[34]. Both CAR-T and CAR-iNKT cells gradually lost IFN-γ production and cytotoxic granule expression with repeated challenges, in line with reports that sustained antigen stimulation progressively impairs T-cell function[35, 36]. Our data indicate that CAR effector dysfunction is associated with a gradual decline in CAR surface expression, likely due to antigen-induced trogocytosis and/or CAR internalization followed by lysosomal degradation[37, 38]. In line with this, we observed trogocytosis and rapid intracellular CAR internalization after antigen contact, followed by partial recycling to the cell surface after a single antigen encounter (data not shown). Actually, repeated antigen exposure has been reported to accelerate CAR internalization and degradation[39].

Finally, our data indicate that the HLA-E/NKG2A inhibitory pathway, which is known to limit cytotoxicity[40–43], promotes tumor escape in a context-dependent manner. This effect was detected in only one of the two GBM cell lines and more pronounced with CAR-iNKT cells, suggesting that immune evasion depends on both tumor-intrinsic heterogeneity[44, 45] and the CAR-effector subset involved.

These escape mechanisms occurred despite the use of a fourth-generation GD2 CAR expressing IL-15, a strategy previously associated with enhanced proliferation and cytotoxicity of CAR- iNKT cells against neuroblastoma[46]. In this context, IL-15 has been shown to support lymphocyte survival and expansion[47], promote IFN-γ and granzyme expression[40], and favor memory-like CD62L- and CD27-expressing effector phenotypes[46, 48, 49]. However, in our study, CAR-IL-15 expression was insufficient to prevent the rapid functional decline of CAR-T and CAR-iNKT cells against glioblastoma cell lines. Moreover, exogenous addition of IL-15 ± IL-7 only partially restored CAR-effector function, mainly by increasing CAR-effector numbers and modulating the NKG2A/NKG2D balance, either by limiting NKG2A or enhancing NKG2D.

Among the strategies evaluated to improve CAR-effector function, IL-12 showed the strongest benefit. Our findings identify IL-12 as a key factor that prevents the progressive dysfunction of CAR-T and CAR-iNKT cells during repeated antigen stimulation. In several tumour models, IL-12 preserves T- and NK-cell cytotoxicity by maintaining effector-cell numbers and sustaining IFN-γ, TNF-α, granzyme, and perforin production through STAT4–T-bet signalling[33, 50–53]. Consistent with these reports, IL-12 maintained cytotoxic activity and effector cytokine production in our model, supporting sustained CAR-effector programming. In particular, the addition of IL-12 induces IFN-γ production by both CAR-T and CAR-iNKT effectors, as demonstrated by intracellular flow cytometry and analysis of culture supernatants in our study. This IL-12-mediated induction of IFN-γ may result from activation of the STAT4 pathway, and may subsequently induce T-bet pathway, as previously described by other groups[33, 51, 54]. Here, we confirm that IFN-γ is a key component of the cytotoxic potential of CAR effectors especially against solid tumors, in T cells[55, 56] and in iNKT cells. IL-12 also promoted an activating NK-receptor profile by enhancing NKG2D signalling, which may help preserve cytotoxicity during sequential stimulation. Although IL-12-induced IFN-γ may upregulate the HLA-E/NKG2A inhibitory axis[29, 40, 43], this was accompanied in our experiments by a predominant increase in NKG2D expression. Future studies should quantify soluble MICA-B during repeated tumour challenges and determine whether IL-12 limits ligand shedding or strengthens activating receptor signalling.

Our results also suggest that IL-12 stabilizes CAR expression at the cell surface during repeated antigen exposure, revealing a previously unrecognized link between cytokine signalling and CAR trafficking. The mechanisms controlling CAR surface persistence remain poorly understood. Continuous antigen engagement can promote CAR internalization and degradation[39], whereas ubiquitination may further limit receptor recycling[57]. We hypothesize that sustained IFN-γ production may contribute to CAR stability by modulating trafficking pathways involving Rab5 and Rab7[58], as well as early and recycling endosomes involving Rab5 and Rab11[38]. Future studies should define the mechanisms that regulate CAR stability by IL-12, an underappreciated determinant of therapeutic efficacy.

Intriguingly, IL-12 preserved CAR-T and CAR-iNKT cell function despite enhancing trogocytosis, a process previously associated with CAR-effector dysfunction[59, 60]. This may reflect IL-12-driven immune synapse formation through LFA-1/ICAM-1 interactions[61, 62], which strengthens integrin-mediated adhesion and promotes microtubule-organizing centre polarization, potentially facilitating trogocytosis. In line with previous studies, increased trogocytosis induced by IL-12 was accompanied by higher exhaustion-marker expression in CAR effectors. Thus, trogocytosis may indicate stronger CAR-effector activation during repeated antigen stimulation, while ultimately contributing to exhaustion. This may explain why, with the fourth-generation CD28 CAR used here, IL-12 extended *in vitro* efficacy by only two additional challenge cycles.

Interestingly, IL-12 also prolonged the cytotoxic activity of T cells transduced with a third-generation 4.1BB.CD28.GD2.CAR, which showed greater *in vitro* cytotoxicity and proliferative capacity than the fourth-generation IL-15-CD28.GD2.CAR-T cells. This advantage may be related to the 4-1BB costimulatory domain, which has been reported to enhance CAR persistence, reduce exhaustion[63], support metabolism and memory-like differentiation[64], and decrease apoptosis[65] compared with CD28-based CARs.

This study has several limitations. First, all experiments were performed *in vitro* and may therefore not fully capture the complexity of TME. Second, GD2.CAR-transduced cells were not purified or sorted after transduction to isolate CAR-expressing cells, resulting in mixed populations in which non-transduced cells may have influenced responses independently of CAR signalling[66].

In summary, our study shows that GD2.CAR-T and GD2.CAR-iNKT cells initially exert potent anti-GBM activity but progressively lose function after repeated antigen exposure. This decline is associated with reduced CAR surface expression, impaired effector activity, GD2 downregulation, trogocytosis, and activation of inhibitory immune pathways, all of which are significantly counteracted by IL-12 supplementation significantly counteracts. These findings support the incorporation of IL-12 signalling into CAR engineering to improve the durability of CAR-based therapies for solid tumors.

## List of abbreviations

GBM: Glioblastoma
CAR: chimeric antigen receptor
PBMCs: peripheral blood mononuclear cells
CNS: central nervous system
TAMs: tumor-associated macrophages
MDSCs: myeloid-derived suppressor cells
TME: tumor microenvironment
FBS: fetal bovine serum
PS: penicillin-streptomycin
MFI: median fluorescence intensity
IL: interleukin
D0: day 0
C-I: cycle 1
C-II: cycle 2
C-III: cycle 3

## Authorship contributions

T.-D.T., C.L., L.G., and J.B. participated in isolating and transducing immune cells with lentiviral and retroviral vectors. T.-D.T., M.-T.R., and C.P. contributed to the study conception and design, supervised data collection and analyses, and interpreted the results. T.-D.T. drafted the manuscript. M.-T.R. and C.P. reviewed the manuscript. T.-D.T., M.-T.R., C.P., G.D., D.M., and L.R. revised the final version. All authors reviewed and approved the final version of the manuscript.

## Acknowledgements

We would like to express our gratitude to the “La Ligue contre le Cancer”, Pr. Gianpietro Dotti’s team (Lineberger Comprehensive Cancer Center, University of North Carolina, Chapel Hill, NC, USA) for generous support of this research.

