## Supplementary Figures for "IL-12 restores the sequential cytotoxic capacities of anti-GD2 CAR-T and CAR-iNKT cells against glioblastoma"

[1] Cell-engineering and immunomodulation of Inflammatory and Neoplastic Disorders (CImIND), UMR 7365 CNRS IMOPA-Biopole, 54500, Vandoeuvre-Les-Nancy, France

[2] Cell Therapy and Tissue Bank Unit, Advanced Therapy Medicinal Products Department, CHRU of Nancy, France

[3] Pediatric onco-hematology department of Nancy University Hospital (CHRU), 54500 Vandoeuvre-Les-Nancy, France

[4] Hematology department of Nancy CHRU, Brabois Hospital, 54500, Vandoeuvre-Les-Nancy, France

[5] Lineberger Comprehensive Cancer Center, University of North Carolina, Chapel Hill, NC, USA

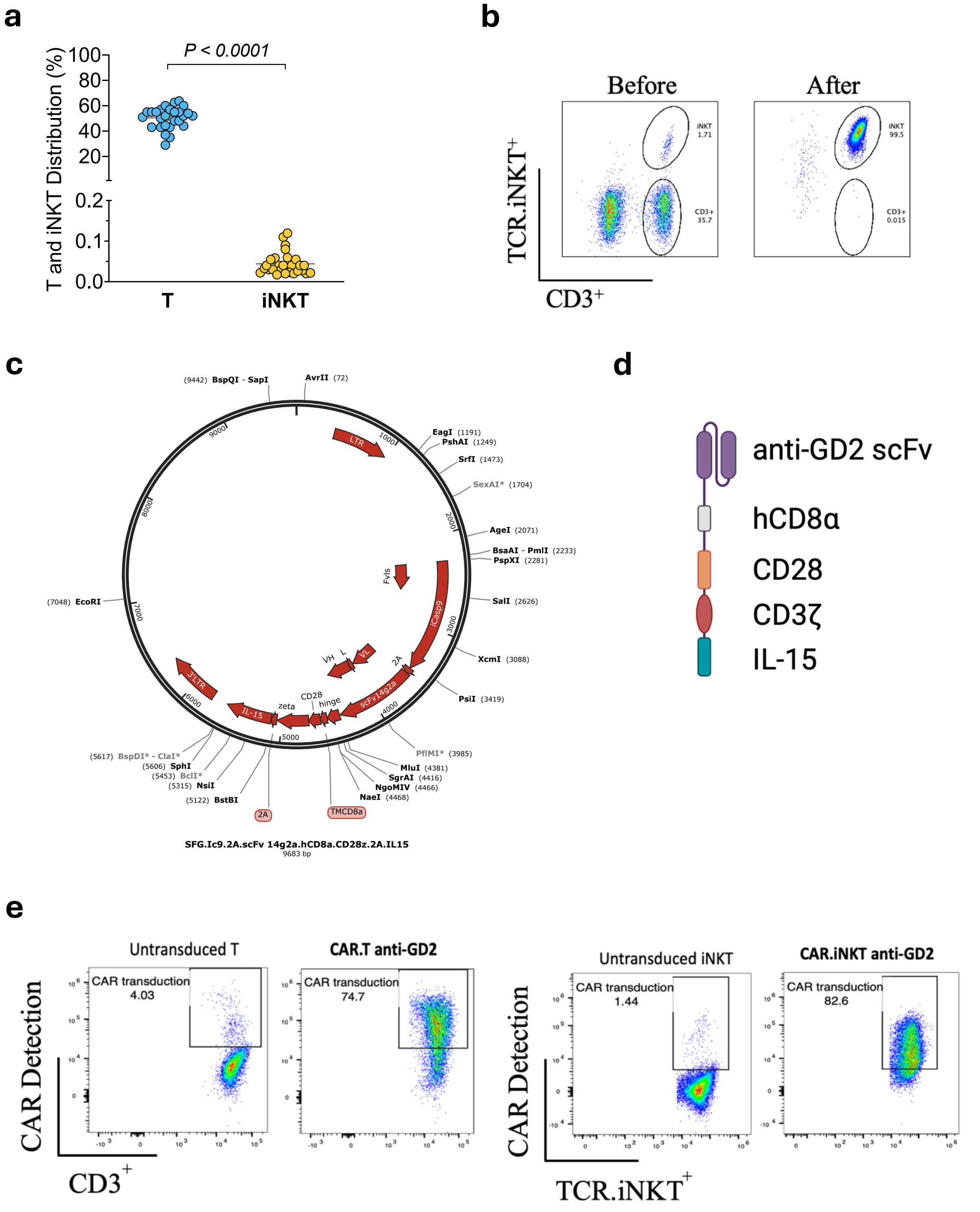


**Figure S1 : Results of isolation and transduction of T and iNKT cells with 4th-generation anti-GD2 CAR retrovirus.**

***a.*** *Distribution of T and iNKT cells before isolation from PBMCs (n=27).* ***b.*** *Percentage of iNKT cells in PBMCs before and after isolation showing over 99% purity.* ***c-d.*** *4th CAR generation anti GD2 retrovirus construct.* ***e.*** *Proportion of CAR expressing effector cells on transduced and untransduced T and iNKT cells by day 11 of in vitro expansion. Comparison performed by Two-way ANOVA test, **** = p < 0.0001.*

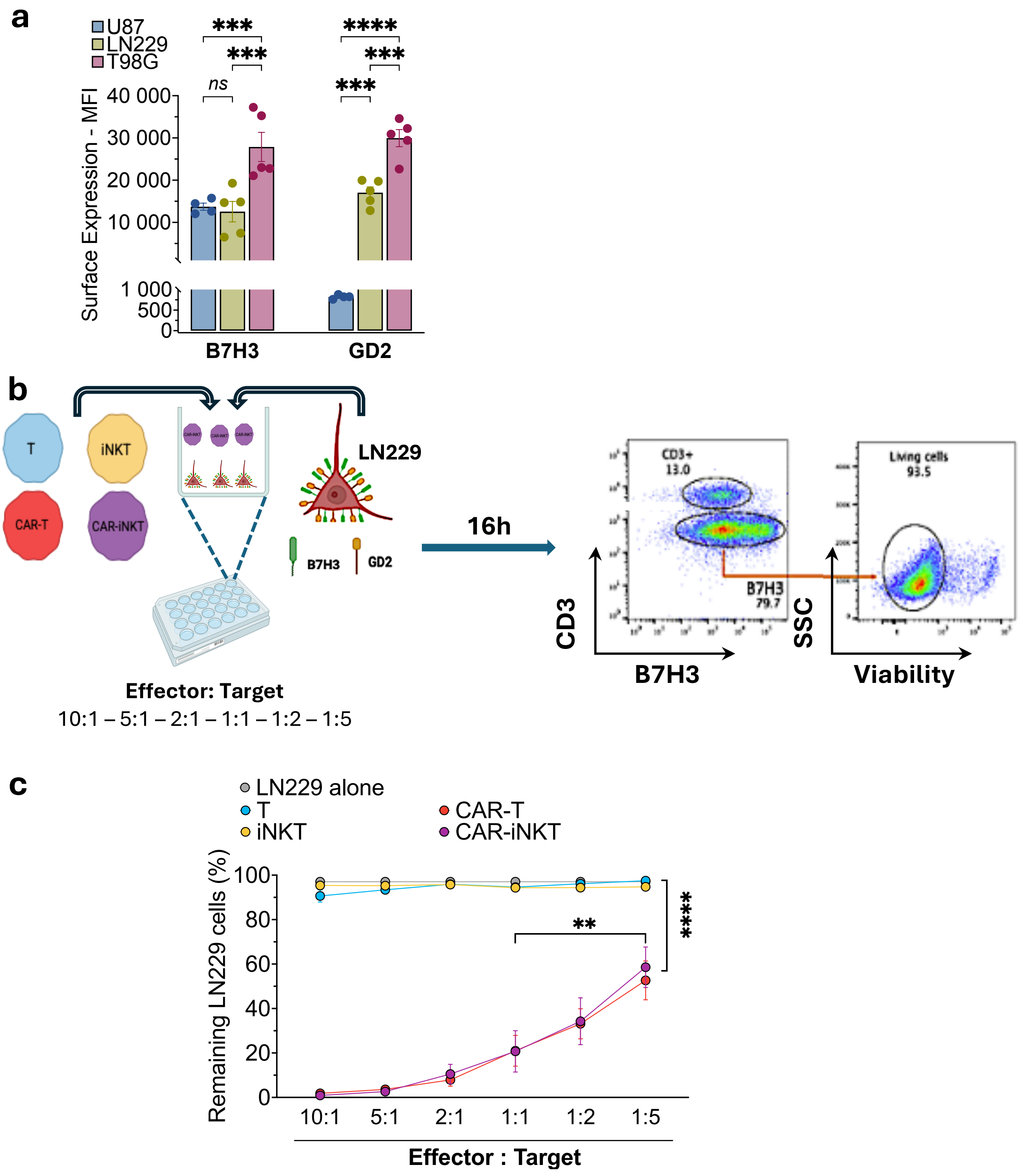


**Figure S2 : Cytotoxicity capacity of CAR effector cells against LN229 at different effector-to-target ratios.**

***a.*** *MFI expression of B7H3 and GD2 was shown, including two GD2-positive cell lines (LN229 and T98G) and one with lower GD2 expression (U87).* ***b.*** *Flowchart of co-cultures including T, iNKT, CAR-T, and CAR-iNKT cells with LN229 at different ratios, E:T – 10 :1, 5 :1, 2 :1, 1 :1, 1 :2, and 1 :5. After 16 hours of co-culture, whole cells in the well were collected and labeled with CD3, B7H3 and viability markers to evaluate the proportions of remaining alive effector ($CD3^+$) and tumor ($B7H3^+$) cells.* ***c.*** *Results of cytotoxic activity of non-transduced cells (T and iNKT cells) and GD2.CAR cells against LN229 after 16 hours are shown as proportions of remaining LN229 cells after the co-cultures. By contrast with untransduced effectors, GD2.CAR-T and CAR-iNKT cells showed similar dose-dependent cytotoxic effects against LN229 cells (n=6). Comparisons were performed with Two-way ANOVA or Tukey's multiple comparisons tests, ns = not significant, ** = $p < 0.01$, *** = $p < 0.001$, **** = $p < 0.0001$.*

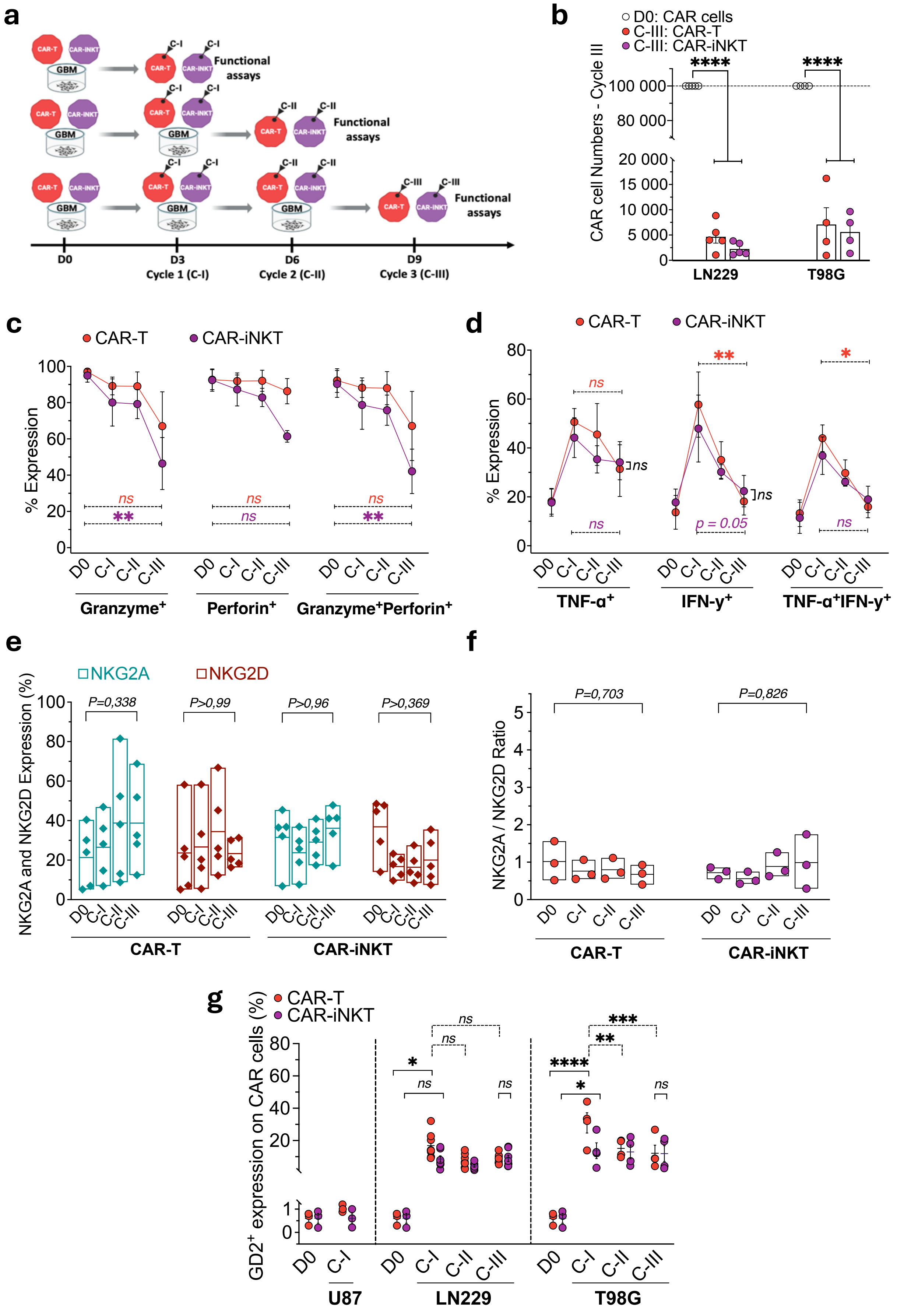
a
CAR-T
CAR-iNKT
GBM
C-I
C-II
C-III
Functional assays
D0
D3 Cycle 1 (C-I)
D6 Cycle 2 (C-II)
D9 Cycle 3 (C-III)
b
D0: CAR cells
C-III: CAR-T
C-III: CAR-iNKT
CAR cell Numbers - Cycle III
LN229
T98G
c
% Expression
Granzyme+
Perforin+
Granzyme+Perforin+
d
TNF-α+
IFN-γ+
TNF-α+IFN-γ+
p = 0.05
e
NKG2A
NKG2D
NKG2A and NKG2D Expression (%)
P=0,338
P>0,99
P>0,96
P>0,369
f
NKG2A / NKG2D Ratio
P=0,703
P=0,826
g
GD2+ expression on CAR cells (%)
U87
LN229
T98G

***Figure S3 : Characterization of CAR-T and CAR-iNKT cells during sequential challenges with GBM cells.***

***a.*** *Flowchart of re-exposure challenges between GBM cells (LN229 and T98G) and GD2.CAR effector cells, beginning with 1 × $10^5$ cells for both GBM and CAR effectors cells. At each 3-day cycle, whole cells from one well in each condition were used for functional assays, while the cells from the remaining wells were seeded with 1 × $10^5$ new tumor cells for another 3-day cycle for a total of 3 cycles.* ***b.*** *Monitoring of CAR effector cell numbers during co-culture with LN229 and T98G cells showing a significant decrease in both GD2.CAR-T and CAR-iNKT cell numbers from 1 × $10^5$ cells at D0 to below 6000 cells after cycle III (n=5, $p < 0.0001$).* ***c-d.*** *Evolution of the proportions of granzyme, perforin TNF-α, and IFN-γ expression by both GD2.CAR-T (red) and CAR-iNKT (purple) cells during challenges against T98G cells.* ***e-f.*** *Proportions of NKG2A and NKG2D expression (**e**), and NKG2A/NKG2D ratio (**f**) on CAR effector cells before (D0) and after each rechallenge cycles against LN229 (**e**) and T98G (**f**).* ***g.*** *Flow cytometric quantification indicated that both CAR-T and CAR-iNKT cells displayed GD2 antigens on their surface following co-culture with $GD2^+$GBM cells and were analysed during the challenge cycles. Trogocytosis was significantly observed at cycle I, but not significantly at the later cycles, although still detected. D0 = day 0, C-I-III = cycles 1-3. Comparisons performed with Two-way ANOVA or Tukey's multiple comparison tests, ns = not significant, * = $p < 0.05$, ** = $p < 0.01$, *** = $p < 0.001$, **** = $p < 0.0001$.*

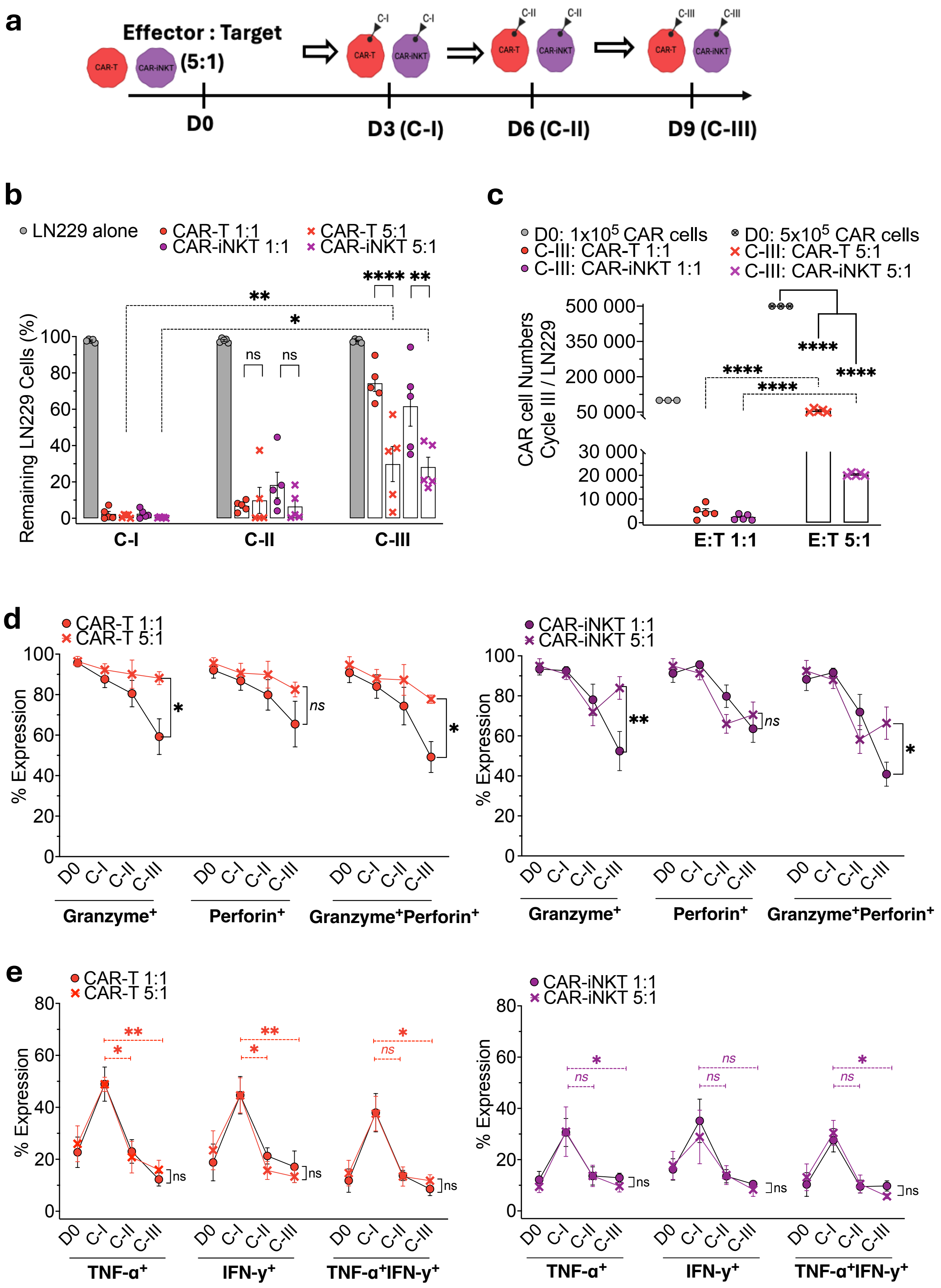
a
Effector : Target
(5:1)
CAR-T
CAR-iNKT
D0
D3 (C-I)
D6 (C-II)
D9 (C-III)
b
LN229 alone
CAR-T 1:1
CAR-iNKT 1:1
CAR-T 5:1
CAR-iNKT 5:1
Remaining LN229 Cells (%)
C-I
C-II
C-III
c
D0: 1x10^5 CAR cells
C-III: CAR-T 1:1
C-III: CAR-iNKT 1:1
D0: 5x10^5 CAR cells
C-III: CAR-T 5:1
C-III: CAR-iNKT 5:1
CAR cell Numbers Cycle III / LN229
E:T 1:1
E:T 5:1
d
% Expression
Granzyme+
Perforin+
Granzyme+Perforin+
e
TNF-α+
IFN-γ+
TNF-α+IFN-γ+

***Figure S4 : Impact of increased effector cell numbers on the functional capacities of CAR-T and CAR-iNKT cells during sequential challenges by LN229.***

***a.*** *Flowchart of re-exposure challenges between LN229 cells and GD2.CAR effector cells beginning with a 5:1 E:T ratio.* ***b.*** *LN229 residual cells after each challenge according to E:T ratio 5:1 (x symbols) and 1:1 (circle symbols). Increased CAR effector cell numbers (5:1 E:T ratio) partially restored tumor cell elimination by both GD2.CAR-T and CAR-iNKT populations by cycle 3 compared to 1:1 ratio (n=5, $p < 0.0001$).* ***c.*** *Monitoring of GD2.CAR effector cells show increased numbers by cycle 3 at 5:1 compared to 1:1 E:T ratios ($p < 0.0001$) but below $5 \times 10^5$ cells put at D0 (($p < 0.0001$).* ***d.*** *Evolution of the proportions of granzyme, perforin or both expression by GD2.CAR-T (red) and CAR-iNKT (purple) cells according to E:T ratios showing a relative maintenance on CAR-T and lower reduction on CAR-iNKT cells by cycle 3.* ***e.*** *Evolution of the proportions of TNF-α, and IFN-γ or both expression on GD2.CAR-T (red) and CAR-iNKT (purple) cells according to E:T ratios showing similar loss of cytokine production in all conditions by cycle 3. D0 = day 0, C-I-III = cycle 1-3. Comparisons performed with Two-way ANOVA, or Tukey's multiple comparisons tests, ns = not significant, * = $p < 0.05$, ** = $p < 0.01$, **** = $p < 0.0001$.*

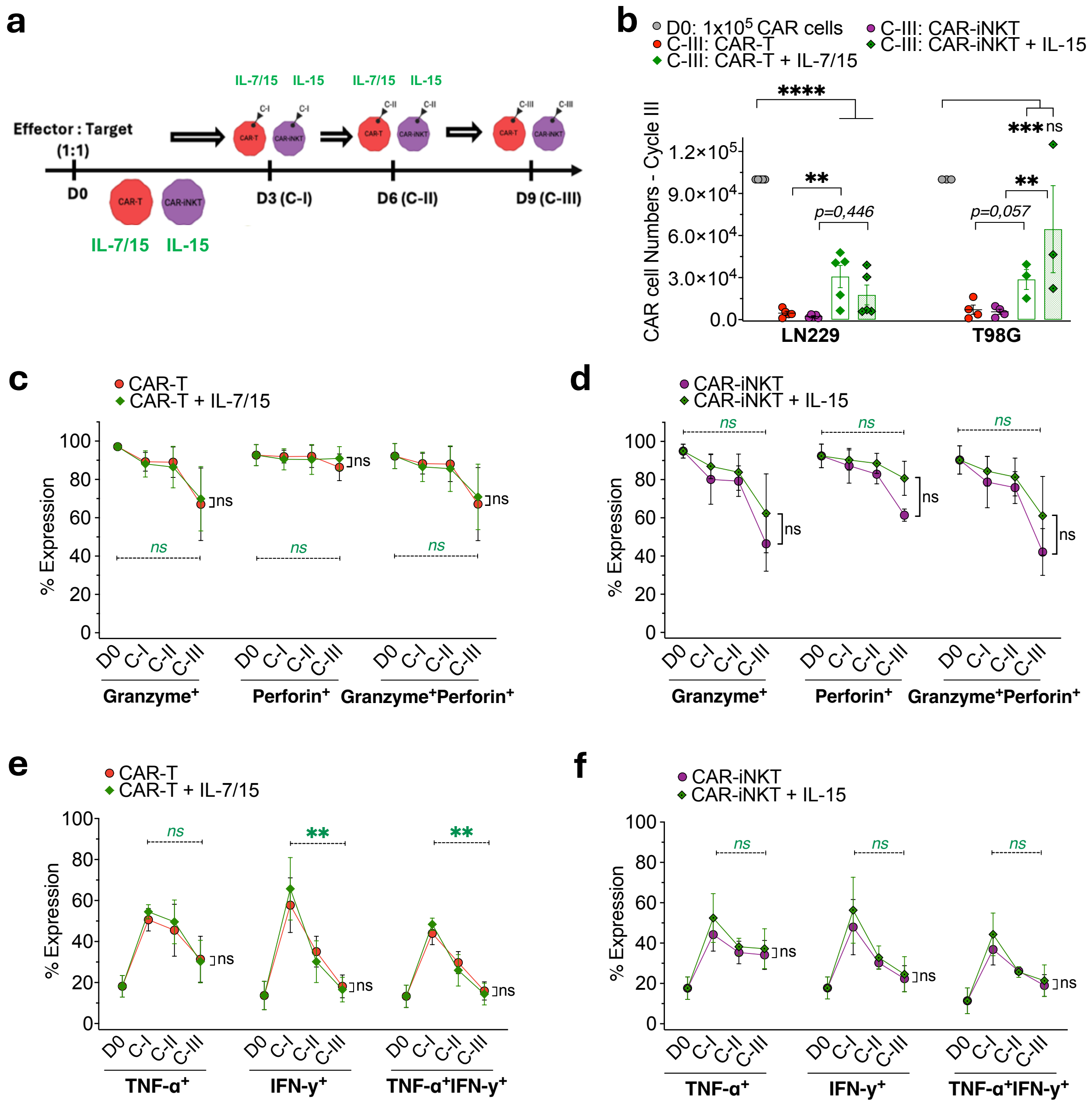


***Figure S5 : Characterization of CAR-T and CAR-iNKT cells in a sequential GBM challenge with the cytokine support.***

*a. Flowchart of re-exposure challenges between LN229 or T98G cells and GD2.CAR effectors at 1:1 ratio with or without cytokine (IL-7/15 for CAR-T and IL-15 for CAR iNKT) or α-GalCer supplementation at each cycle. **b.** Monitoring of GD2.CAR effector cell numbers by cycle 3 show significant increase of CAR-T cells with IL-7 and IL-15 supplementation (n=3-5, $p < 0.001$) against LN229 and of CAR-iNKT cells (n=3-5, $p<0.001$) with IL15 supplementation against T98G in comparison to D0. However, numbers of remaining effectors by cycle 3 were below $1 \times 10^5$ cells put at D0 in all conditions ($p < 0.05$). **c-d.** Similar loss of proportions of granzyme, perforin or both expression by GD2.CAR-T (left) and CAR-iNKT (right) cells with or without cytokine supplementation over cycles against T98G. **e-f.** Similar loss of proportions of TNF-α, and IFN-γ or both expression on GD2.CAR-T (left) and CAR-iNKT (right) cells with or without cytokine supplementation over cycles against T98G. D0 = day 0, C-I-III = cycle 1-3. Comparisons performed with Two-way ANOVA, or Tukey's multiple comparisons tests, ns = not significant, * = $p < 0.05$, ** = $p < 0.01$, **** = $p < 0.0001$.*

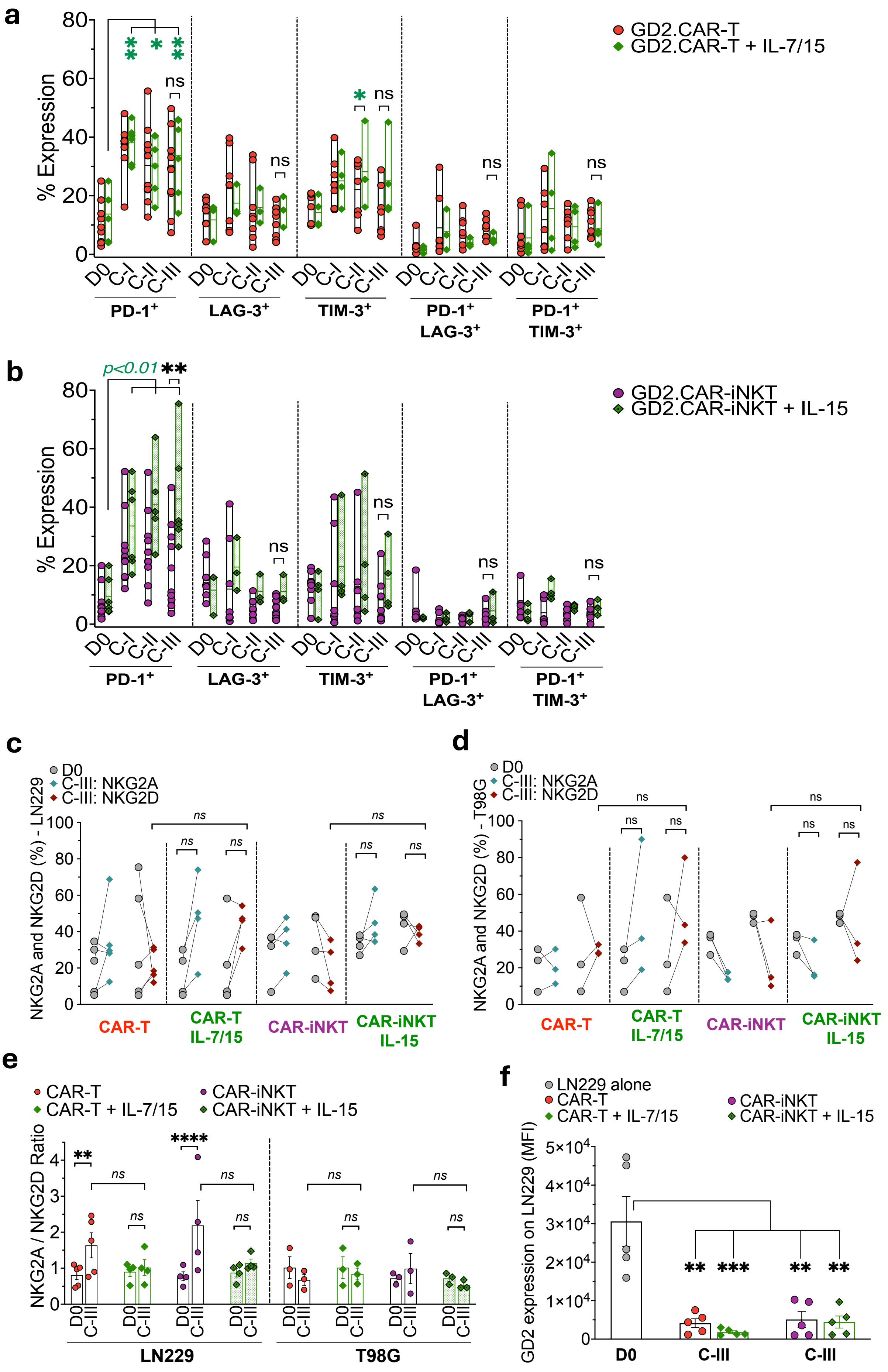
a
GD2.CAR-T
GD2.CAR-T + IL-7/15
% Expression
D0 C-I C-II C-III
PD-1+
LAG-3+
TIM-3+
PD-1+ LAG-3+
PD-1+ TIM-3+
ns
b
p<0.01 **
GD2.CAR-iNKT
GD2.CAR-iNKT + IL-15
% Expression
c
D0
C-III: NKG2A
C-III: NKG2D
NKG2A and NKG2D (%) - LN229
CAR-T
CAR-T IL-7/15
CAR-iNKT
CAR-iNKT IL-15
d
NKG2A and NKG2D (%) - T98G
e
CAR-T
CAR-T + IL-7/15
CAR-iNKT
CAR-iNKT + IL-15
NKG2A / NKG2D Ratio
LN229
T98G
f
LN229 alone
GD2 expression on LN229 (MFI)
D0
C-III

***Figure S6 : Impact of cytokine supplementation on exhaustion markers and NK receptor expression on CAR-T and CAR-iNKT cells across sequential GBM challenge along with GD2 expression on LN229 cells.***

***a-b.** Similar expression of exhaustion markers PD-1, LAG-3, TIM-3 alone or in combination to PD-1 on GD2 CAR-T (**a**) and CAR-iNKT (**b**) cells with or without cytokine supplementation during LN229 challenge cycles (n=3-6). **c-d.** Expression of NKG2A and NKG2D on GD2.CAR-T and GD2.CAR-iNKT cells before and after LN229 (**c**) and T98G (**d**) sequential challenges with or without cytokine supplementation showing a non-significant increase of NKG2D expression on both GD2 CAR T and CAR-iNKT cells challenged by LN229 (n=3-4). **e.** NKG2A/NKG2D ratio increase observed with LN229 on CAR effector cells by cycle 3 was did not seen in the presence of cytokine supplementation. **f.** Median Fluorescence Intensity (MFI) of GD2 antigen on LN229 cells measured by cycle 3, at the time of tumor escape to CAR-T and CAR-iNKT cells showed similar antigen expression loss with or without cytokine supplementation (n=5, p < 0.01 – 0.001). D0 = day 0, C-I-III = cycle 1-3. Comparisons performed with Two-way ANOVA, or Tukey's multiple comparisons tests, ns = not significant, * = p < 0.05, ** = p < 0.01, *** = p < 0.001, **** = p < 0.0001.*

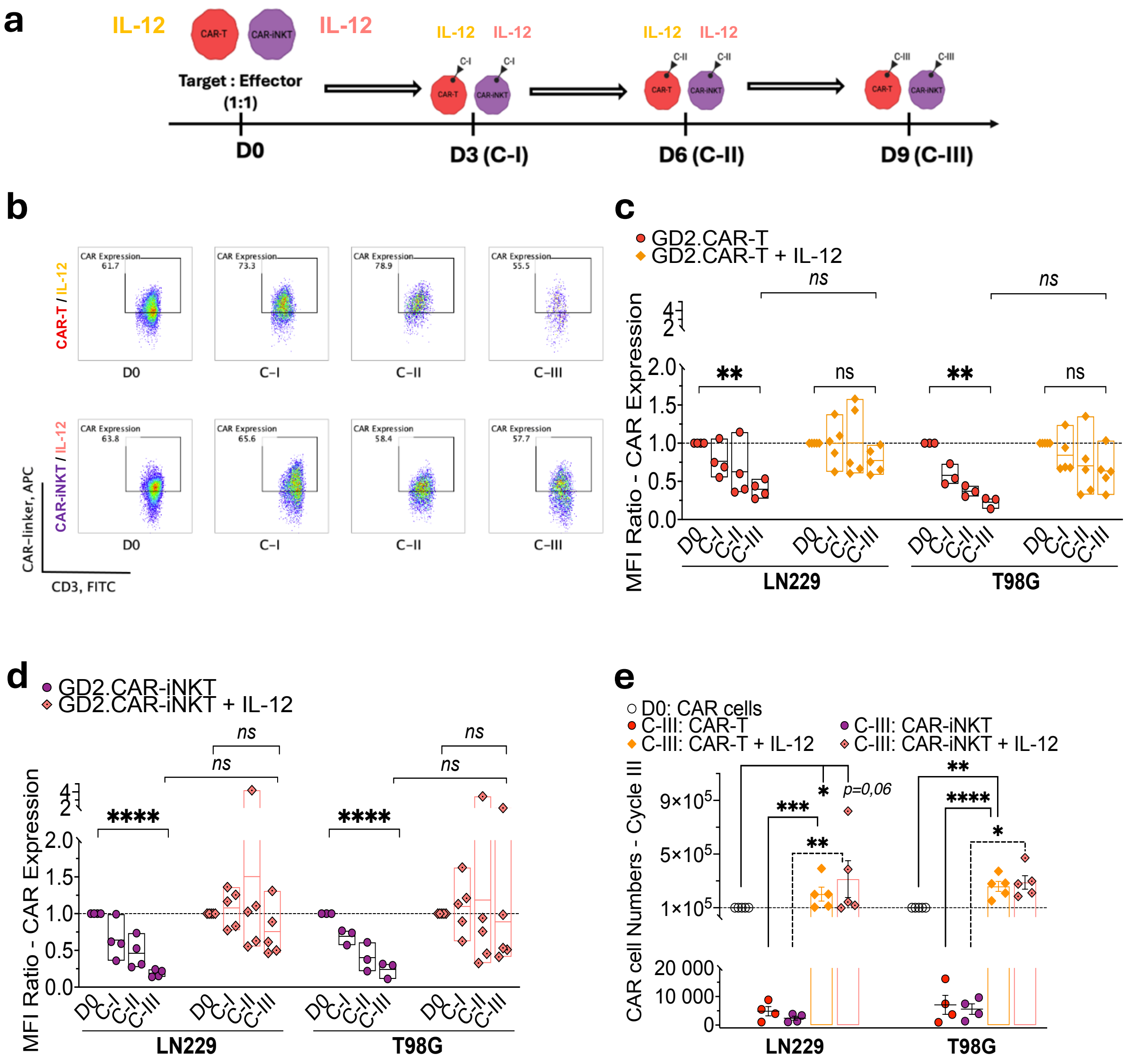


***Figure S7 : Impact of IL-12 on CAR expression and effector cell numbers across GBM challenge cycles.***

***a.*** *Flowchart of re-exposure challenges between GBM cells and GD2.CAR effector cells at a 1:1 ratio with or without IL-12 addition during all cycles.* ***b.*** *Flow cytometry plots showing stable CAR expression at the surface of GD2 CAR-T and CAR-iNKT cells across GBM challenge cycles in the presence of IL-12 addition.* ***c-d.*** *Median Fluorescence Intensity (MFI) of CAR molecule on GD2 CAR-effector cells reported as a ratio relative to D0 MFI was calculated across 3 cycles on CAR-T (**c**) and CAR-iNKT cells (**d**) with or without IL-12 (n=4-5). The addition of IL-12 was associated with the stability of CAR expression by contrast with progressive CAR expression loss over cycles on both effectors in the absence of IL-12.* ***e.*** *GD2 CAR effector cell numbers by cycle 3 for both CAR-T and CAR-iNKT cells with or without IL-12 addition showing a significant expansion of both CAR effectors in comparison to D0 and to cycle 3 without IL-12 (n=4-5, $p<0.001$). D0 = day 0, C-I-III = cycle 1-3. Comparisons performed with Two-way ANOVA, or Tukey's multiple comparisons tests, ns = not significant, * = $p < 0.05$, ** = $p < 0.01$, *** = $p < 0.001$, **** = $p < 0.0001$.*

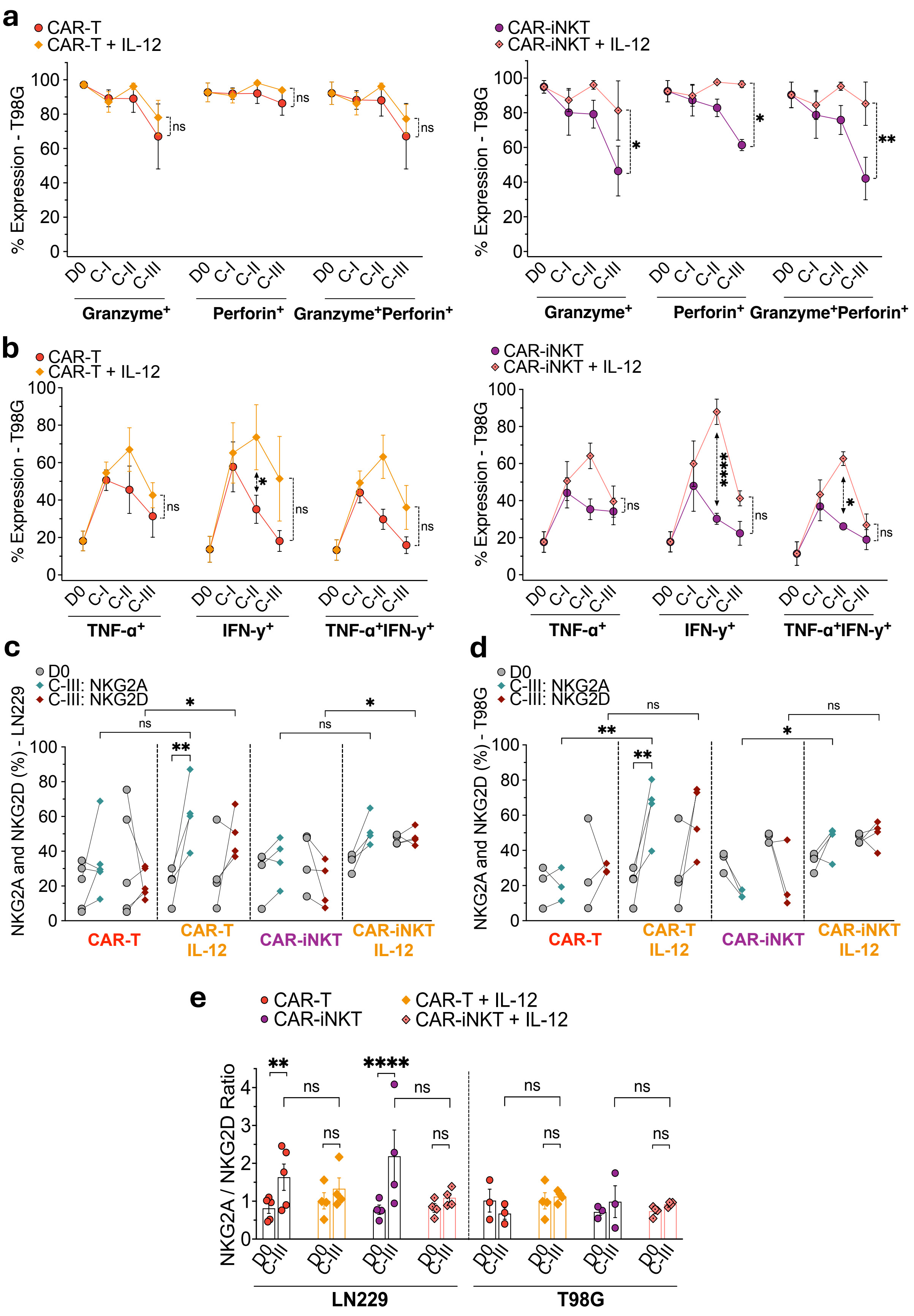
a
CAR-T
CAR-T + IL-12
% Expression - T98G
Granzyme+
Perforin+
Granzyme+Perforin+
CAR-iNKT
CAR-iNKT + IL-12
b
TNF-α+
IFN-γ+
TNF-α+IFN-γ+
c
D0
C-III: NKG2A
C-III: NKG2D
NKG2A and NKG2D (%) - LN229
CAR-T
CAR-T IL-12
CAR-iNKT
CAR-iNKT IL-12
d
NKG2A and NKG2D (%) - T98G
e
NKG2A / NKG2D Ratio
LN229
T98G

***Figure S8 : Impact of IL-12 on GD2.CAR-T and CAR-iNKT cell cytokine production, cytolytic molecule expression and NKG2A/NKG2D ratio across T98G challenge cycles.***

***a.*** *Proportions of granzyme, perforin or both expression on GD2.CAR effectors during T98G challenges were significantly increased on CAR-iNKT (right) by cyle 3 when IL-12 was added across cycles (n=3, p<0.001 versus no IL-12).* ***b.*** *Proportions of GD2.CAR T and CAR.iNKT cells producing FN-gamma were significantly at the end of cycle 2 of T98G rechallenge in the presence of IL-12 (n=3, p<0.05 for CAR-T and <0.001 for CAR-iNKT).* ***c-d.*** *Expression of NKG2A and NKG2D on GD2.CAR-T and GD2.CAR-iNKT cells before and after LN229 (**c**) and T98G (**d**) sequential challenges with or without IL-12 addition show a significant increase of NKG2D expression on both GD2 CAR T and CAR-iNKT cells challenged by LN229 after cycle 3 (n=3-4, p<0.05 versus no IL-12), and an increase of NKG2A on both CAR-effectors by cycle 3 with T98G rechallenge (n=3-4, p<0.05 versus no IL-12).* ***e.*** *NKG2A/NKG2D ratio increase observed with LN229 on CAR effector cells by cycle 3 was not observed in the presence of cytokine supplementation. D0 = day 0, C-I-III = cycle 1-3. Comparisons performed with Two-way ANOVA, or Tukey's multiple comparisons tests, ns = not significant, * = p < 0.05, ** = p < 0.01, *** = p < 0.001, **** = p < 0.0001.*

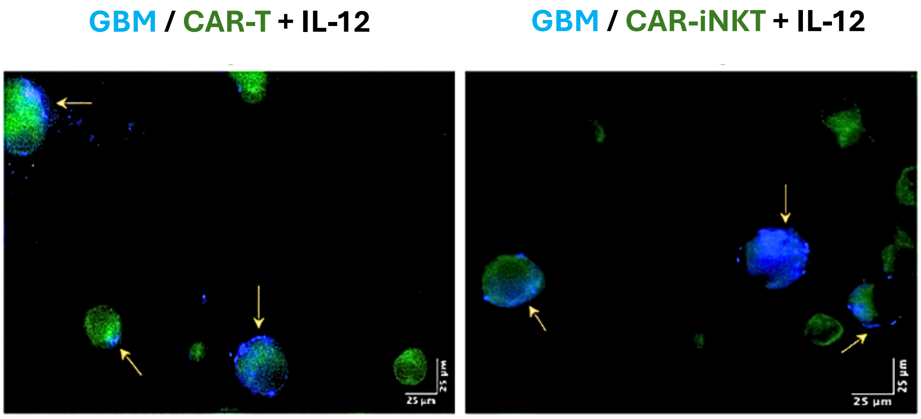


***Figure S9 : Impact of IL-12 on Trogocytosis of GD2 by CAR-T and CAR-iNKT cells***

*Fluorescence microscopy images (40x objectives) show GD2 antigen and CD3 staining on CAR-T and CAR-iNKT cells in the presence of IL-12 supplementation. CAR cells were collected after 4 hours. Yellow arrows indicate the presence of GD2 antigens (blue fluorescence, GD2-BV421) on the surface of effector cells (green fluorescence, CD3-FITC).*

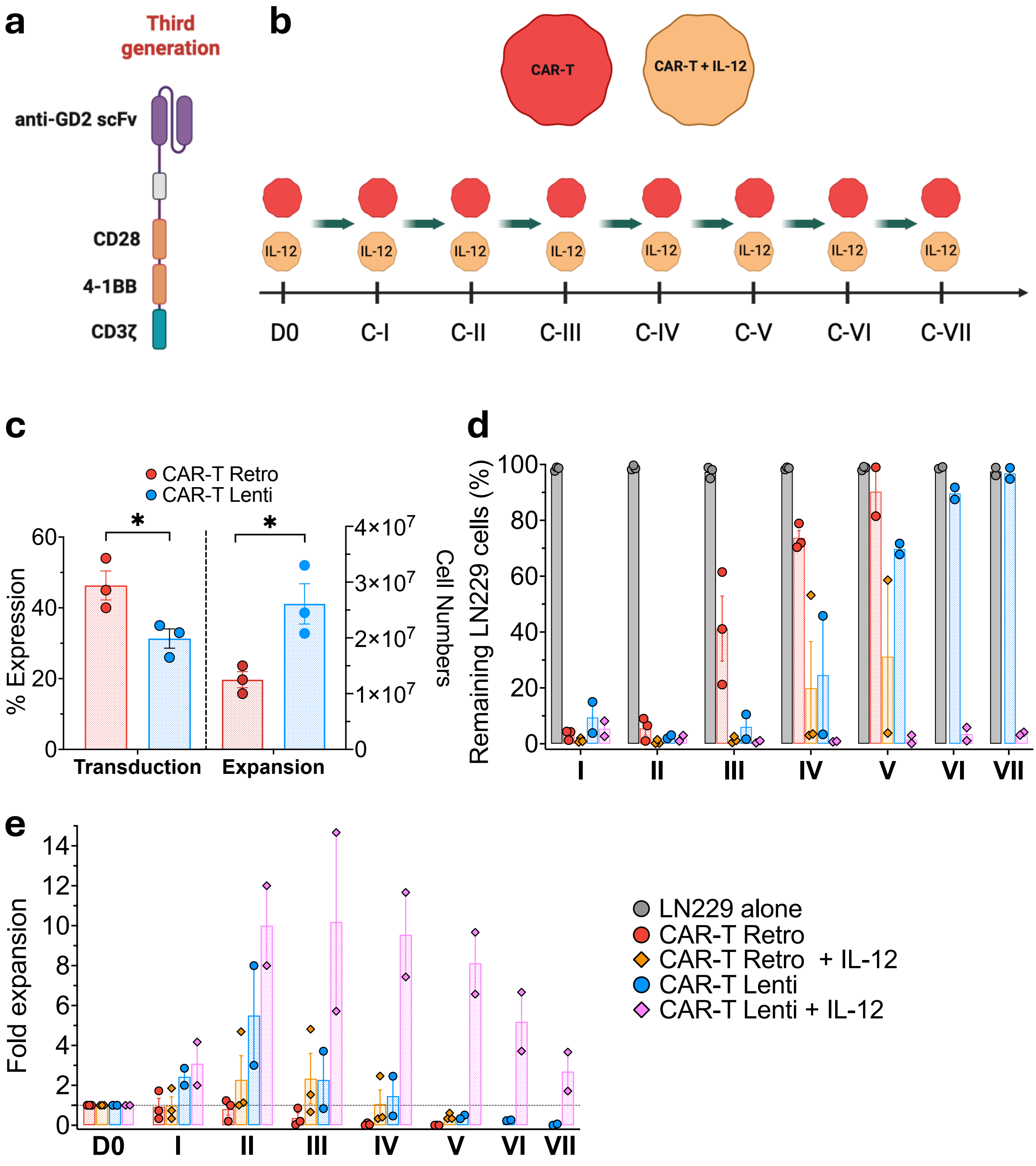


***Figure S10 : IL-12 enhances the anti-tumor activity of CAR-T cells transduced with a third-generation anti-GD2.CAR.***

***a.** Anti-GD2.3$^{rd}$ generation CAR construction containing two costimulatory domains (4.1BB and CD28) inserted in a lentivirus. **b.** Flowchart illustrating the number of tumor challenges performed with the two anti-GD2 CAR-T cells produced (4$^{th}$ generation retrovirus and 3$^{rd}$ generation lentivirus) supplemented with or without IL-12 during each 3 days challenge cycle. **c.** Transduction efficacy and CAR-T numbers obtained 14 days after transduction of $10^5$ isolated T cells with a 4th generation (retrovirus) or a 3rd generation (lentivirus) anti-GD2 CAR (n=3 for each). **d.** Percentage of residual LN229 cells after co-culture with retro-GD2.CAR-T alone (red) or with IL12 (orange) (n=3) and with lenti-GD2.CAR-T alone (blue) or with IL-12 (pink) (n=2) after each tumor challenge cycle. **e.** Fold expansion of CAR-T cells relative to D0 after each LN229 challenge cycle for retro-GD2.CAR-T alone (red) or with IL12 (orange) (n=3) and with lenti-GD2.CAR-T alone (blue) or with IL-12 (pink) (n=2).*
